# Live-cell imaging and lipidomics of low density lipoprotein containing intrinsically fluorescent cholesteryl esters

**DOI:** 10.1101/2025.08.05.668602

**Authors:** Alice Dupont Juhl, Richard R. Sprenger, Maciej Modzel, Senjuti Halder, Maria Szomek, Xuanjiang Xu, Christer S. Ejsing, Douglas F. Covey, Daniel Wüstner

## Abstract

Low density lipoprotein (LDL) delivers cholesterol to cells in the body in the form of cholesteryl esters (CEs), and dysfunction of this pathway is associated with various diseases. Due to the lack of suitable tools, our understanding of the intracellular transport and hydrolysis of CEs is limited. We present a novel approach for studying LDL-derived CEs in cells, using fatty acyl chain conjugates of the intrinsically fluorescent cholestatrienol (CTL). We demonstrate that CTL esters reconstituted into LDL particles are hydrolyzed in late endosomes and lysosomes (LE/LYSs) by acid lipase, while an LDL-derived CTL ether analog cannot leave LE/LYSs. Using live-cell imaging, lipidomics, and multimodal Bayesian modeling, we discover a sequential biphasic transport of LDL-derived CTL to LE/LYSs matching the kinetics of the lysosomal hydrolysis of LDL-associated CTL-ester with a half-time of 3.0 hours. Hydrolyzed CTL derived from LDL-associated CTL esters is rapidly re-esterified with a similar half-time and stored in lipid droplets, demonstrating efficient sterol transport to the endoplasmic reticulum (ER). The latter is supported by the detection of a faint CTL staining in the ER and by extensive contact formation between endo-lysosomes containing LDL and ER tubules. Using lipidomics and Bayesian kinetic modeling, we also track LDL-derived CEs and triacylglycerols in cells and determine the uptake kinetics for each lipid species individually. Our novel approach allows for precise measurement of post-endocytic trafficking and metabolism of LDL-derived cholesterol and other lipids in living cells.

## Introduction

Mammalian cells have a regulated homeostasis of cholesterol with *de novo* synthesis and cellular uptake controlled by transcriptional processes located primarily in the endoplasmic reticulum (ER). The abundance of cholesterol in the ER is low under normal conditions, and dedicated mechanisms exist in the ER to maintain cholesterol homeostasis. Excess cholesterol is esterified by sterol O-acyltransferases (i.e., SOAT1, SOAT2), also named acyl-CoA acyl transferase (ACAT), and stored as cholesteryl esters (CEs) in lipid droplets [1, 2]. A drop below about 5 mol% signals cellular cholesterol depletion to the cholesterol-sensing SCAP protein and triggers transcription of cholesterol target genes via the sterol response element binding protein (SREBP) [3, 4]. These genes code for enzymes of the biosynthetic pathway as well as the receptor for low density lipoprotein (LDL-R), which mediates uptake of cholesteryl ester-rich LDL particles by clathrin-mediated endocytosis (CME) [4–7]. Internalized LDL dissociates from its receptor in the endosomes due to acidification of these organelles via the vacuolar proton pump [8, 9]. The LDL particles released from their receptor in early endosomes are transported to late endosomes and lysosomes (LE/LYSs), where their degradation starts after a lag phase of about 1-3h post internalization [10–13]. Biochemical studies using radioactive tracers showed that lysosomal acid lipase (LAL, i.e., lysosomal integral protein A, LIPA) mediates the hydrolysis of the majority of the LDL-associated CEs [13–18].

The trafficking itineraries and metabolic fates of LDL-derived cholesterol, once CEs are hydrolyzed, are less-well defined [19, 20]. By reconstituting LDL particles with radioactive CEs, Neufeld and coworkers provided evidence for a direct trafficking route of the liberated cholesterol from LE/LYSs to the plasma membrane (PM) [21]. They suggested that this route comprises about 70% of the LDL-derived cholesterol, while the remaining 30% traffics to the ER. Similar conclusions were drawn by Liscum and co-workers, who employed small molecule inhibitors of lysosomal cholesterol export, such as U18666A and imipramine, to show that a portion of LDL-derived cholesterol directly traffics to the ER [22]. Others have argued that the entire pool of liberated LDL-derived cholesterol first passes through the PM before reaching the ER [23, 24]. Dysregulated uptake of cholesterol from modified LDL particles takes place during the development of atherosclerosis, which leads to massive re-esterification of excess cholesterol by ACAT1 and foam cell formation in macrophages [19, 25, 26]. In neurodegenerative diseases, such as Wolman and Niemann Pick type C disease, impaired hydrolysis and/or transport of the LDL-derived cholesterol out of the lysosomal compartment leads to lysosome dysfunction [27, 28]. LDL-derived cholesterol also plays an important role in the development of brain cancers, as glioblastoma cells allegedly shut down their endogenous production of cholesterol and rely primarily on imported LDL cholesteryl esters for growth and survival [29]. Thus, understanding the mechanisms underlying cellular processing of LDL-derived cholesterol is instrumental for developing novel therapies against these devastating diseases.

Our current understanding of intracellular transport and metabolism of LDL-derived cholesterol relies mainly on reconstituting radioactively labeled CEs into LDL and following their transport to subcellular organelles by membrane fractionation and scintillation counting. While being powerful, this approach suffers from impurities arising from organelle purification, limited time resolution, and the fact that no directional information is available. Also, subcellular organelles of many cells are pooled before lipid analysis, preventing any insight into cell-to-cell heterogeneity of lipoprotein transport. Classical fluorescence imaging-based approaches use a combination of targeted gene knockouts and fluorescence staining with the sterol-binding polyene filipin or cholesterol sensing proteins, like AloD4, perifringolysin or the GRAMD1 protein domains, to obtain insight into the spatial distribution of various cholesterol pools [30–32]. These tools were employed to identify a variety of proteins, whose genetic ablation causes cholesterol accumulation in LE/LYSs and/or impaired delivery of LDL-derived cholesterol to ACAT in the ER, including members of the oxysterol binding protein family [33–37], cytosolic or ER-bound sterol transfer proteins containing the START domain [38–40] as well as ER-resident transporters containing a sterol-binding GRAM-domain [41–43]. However, given the complexity of sterol trafficking pathways, cholesterol buildup in a given compartment or lack thereof does not necessarily translate into an understanding of the sterol transport dynamics or trafficking routes between individual organelles. For example, increased filipin staining in a subcellular compartment after loading cells with LDL could be the result of not only cholesterol accumulation from the LDL source but also a result of the buildup of endogenous cholesterol. Since filipin is not specific for cholesterol, accumulation of other sterols, such as oxysterols cannot be distinguished by filipin staining either. Furthermore, the kinetics of sterol trafficking between organelles cannot be determined, and the actual trafficking route of the LDL-derived cholesterol is thereby overlooked by this standard approach.

Here, we introduce a novel method to directly follow the intracellular trafficking and metabolic processing of LDL-derived cholesterol using minimally modified fluorescent analogs of CEs. We chemically link cholestatrienol (CTL), differing from cholesterol only in having two additional double bonds in the ring system (Figure 1A), to the fatty acid oleic acid either as a hydrolysable ester (CTL-ester) or non-hydrolysable ether (CTL-ether). We show by live-cell imaging combined with lipid mass spectrometry (Lipid-MS) that these probes reconstituted into LDL particles have different fates in human fibroblasts; while the unnatural CTL-ether accumulates in LE/LYSs, the natural CTL-ester is hydrolyzed by LAL activity in LE/LYSs. Already after 6h, some free CTL was re-esterified, primarily with an octadecanoyl chain to stearic acid, while we also found esterification to polyunsaturated acyl chains after 24h by Lipid-MS. By implementing a hierarchical, multi-modal Bayesian model, we show that both, the increase in endo-lysosomal fluorescence of LDL-derived CTL as well as the hydrolysis of CTL-esters detected by Lipid-MS can be described by a two-step sequential kinetic process with shared rate constants. Using the same modeling approach, we also quantify the kinetics of intracellular accumulation of LDL-derived CEs and TAGs. Finally, we observed and quantified close appositions of LE/LYSs containing LDL with the ER, which could provide a roadway for lipid transfer between both compartments. Our results establish a novel toolbox for direct visualization and quantitative analysis of trafficking and metabolic conversion of lipoprotein-derived lipids in living cells. Our new methods should find wide application for understanding cholesterol homeostatic pathways and disease pathogenesis.

**Figure 1.**
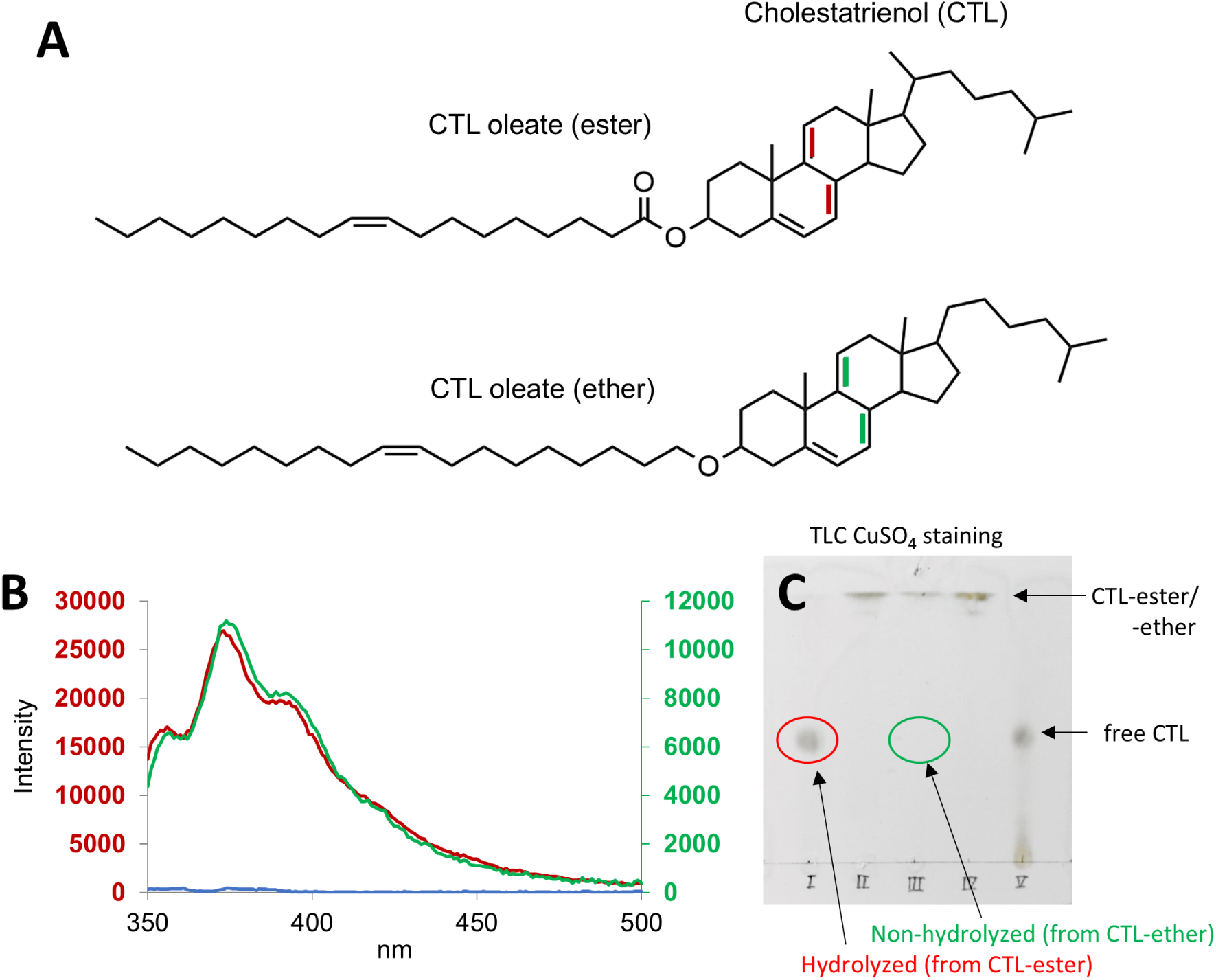
Structure and properties of intrinsically fluorescent CTL 18:1 ester and ether. A, cholestatrienol (CTL) oleate ester and ether are shown with differences to the corresponding cholesterol ester and ether being two additional double bonds in the steroid ring system (indicated in red for the ester and in green for the ether). B, fluorescence emission spectra for 0.2 mg/ml LDL reconstituted with either CTL-oleate ester (red line and left axis) or CTL-oleate ether (green line and right axis), excited at 328 nm. The spectrum of the same amount of unlabeled LDL is shown for comparison (blue). C, thin layer chromatography (TLC) of CTL-ester and ether after treatment with cholesteryl esterase and development of silica plates with copper sulfate. Red circle indicates hydrolyzed CTL obtained from the CTL-ester. The same position is empty for the CTL-ether (green circle). See the main text for further details.

## RESULTS AND DISCUSSION

### Binding and endocytosis of LDL containing fluorescent cholesteryl esters and ethers

Our CTL 18:1-ester and -ether probes contain identical conjugated double bonds in the steroid ring system and differ only in the linkage of the C18:1 hydrocarbon chain to the sterol backbone (Figure 1A). Accordingly, upon reconstitution into LDL (see Figure S1A), their fluorescence spectra are very similar (Figure 1B). The reconstituted particles contain more than 90% sterols, mostly CEs and some cholesterol, with the remaining lipids being comprised of glycerophospholipids, di-and triacylglycerols as well as sphingolipids (Figure S1B). CTL- esters reconstituted into LDL must be tightly packed in the core of the lipoprotein particles, as adding Triton X-100 caused a slight increase in CTL fluorescence as a result of dequenching upon solubilization of the particles (Figure S1C). The main glycerophospholipid in LDL was phosphatidylcholine (PC) followed by various species of phosphatidylethanolamine (PE, Tab. S1). Reconstituted LDL contained additionally sphingomyelin (SM) and small amounts of ceramides (Tab. S2). Together, the composition of the reconstituted LDL particles resembles that of native LDL and is in agreement with previous studies [44].

The fluorescent ester, but not the CTL-ether, can be hydrolyzed by a neutral lipase, as inferred from thin layer chromatography (TLC) plates (Figure 1C). Incubating human fibroblasts with LDL containing CTL-ester overnight resulted in strong fluorescence in the PM and intracellular vesicles (Figure 2A). When excess unlabeled LDL was added during this incubation, the cell-associated CTL fluorescence was greatly reduced (Figure 2A-D). Similar results were found for LDL containing CTL-ether (Figure S2A and B), indicating that the nature of the sterol esters does not affect endocytosis of the LDL particles. This is in line with earlier findings, showing that the apoprotein portion in LDL controls the cellular uptake of the lipoprotein [45]. Remaining cell-associated fluorescence of LDL containing either CTL-esters or -ethers is likely due to binding of the particles to other receptors than the LDL-receptor, as has been observed already in the seminal studies by Goldstein and Brown (1974 and 1975) [13, 15]. These particles could be internalized by an endocytic pathway independent of the LDL-receptor, which is not saturable by excess LDL. Indeed, such a second uptake pathway has been proposed already in these original studies and later identified as being mediated by lipoprotein lipase (LPL) binding to proteoglycans on the surface of fibroblasts and other cells [46, 47]. We suggest that such a pathway also contributes to the uptake and lysosomal accumulation of some LDL containing CTL-esters in our experiments.

**Figure 2.**
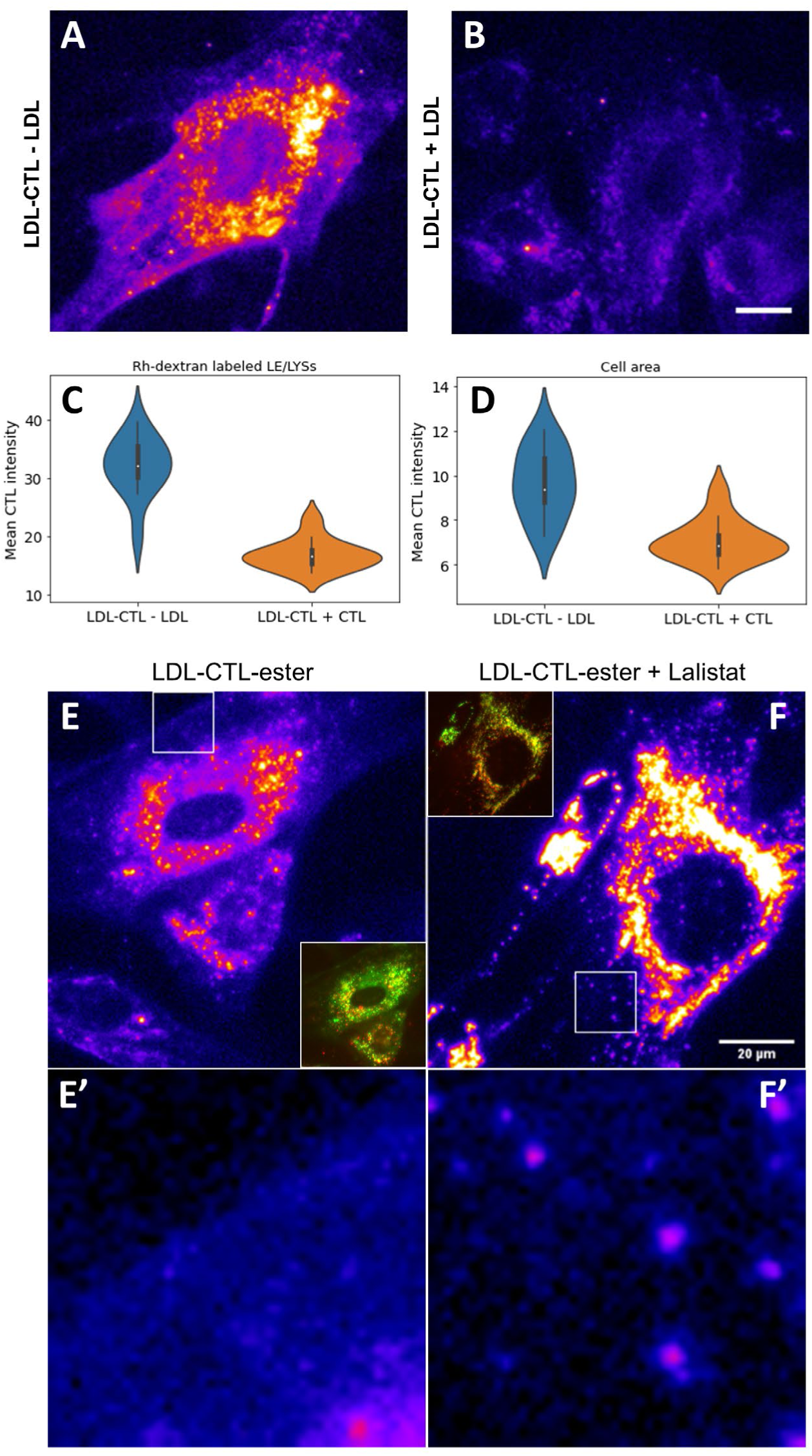
Uptake and lysosomal processing of LDL containing CTL oleate ester. A-C, human fibroblasts were incubated in LPDS medium containing 0.1 mg/ml LDL with CTL-oleate ester and 0.5 mg/ml Rh-dextran to co-stain LE/LYSs overnight, either in the absence (‘-LDL’, right panel in A) or presence of 4 mg/ml unlabeled LDL (‘+ LDL’, left panel in A). CTL fluorescence was quantified in endolysosomes based on the intensity of Rh-dextran (C) or as cell-associated CTL fluorescence (D) for cells incubated with (blue bars) or without excess LDL. The data is shown as mean intensity ± S.D. of the pooled intensities from two independent experiments. E, F, cells were incubated in the absence (E, E’) or presence of the LAL inhibitor Lalistat (F, F’, final concentration 6 µM) for 6h, followed by loading with Rh-dextran and labeled LDL overnight, as described above. For the experiment with Lalistat (F, F’), the inhibitor was present during the incubation with labeled LDL. Inset in E and F shows co-localization of CTL(ester) fluorescence (pseudo-colored in green) with Rh-dextran (red). E’ and F’ are zoomed versions of the rectangular white box in E and F, respectively. They show an enlarged version of the PM with prominent CTL fluorescence in the absence (E’) but not in the presence of Lalistat (F’).

Hydrolysis of CEs derived from LDL has been shown to take place in LE/LYSs primarily by acid lipase, and to test whether this also applies to our novel fluorescent sterol esters, we incubated fibroblasts with LDL containing CTL-ester in the presence of the acid lipase inhibitor Lalistat (Figure 2E and F). We found that in the presence of Lalistat, lysosomal staining of CTL-ester is strongly increased, while that of the PM is very low compared to cells incubated in the absence of Lalistat. Lysosomal targeting was confirmed by co-staining cells with rhodamine-dextran (Rh-dextran; Figure 2E and F, inset). Since the fluorescence of CTL-ester is indistinguishable from that of free CTL, we can only indirectly infer that the PM staining observed for non-treated cells stems from free CTL, as such staining was much lower in the presence of Lalistat (Figure 2E’ and F’). Strong endo-lysosomal fluorescence of the sterol was also found in fibroblasts incubated with LDL containing CTL-ether (Figure S2B). Note also that fluorescence of CTL in the PM of cells after 24h incubation with LDL containing CTL-ester in the absence of the lipase inhibitor is homogeneous, in stark contrast to the patchy PM staining observed in cells incubated with LDL containing CTL-ether at the same time point (Figure S2C). We did not detect any free CTL by Lipid-MS when loading cells with LDL containing CTL-ether, confirming that the CTL 18:1-ether cannot be hydrolyzed in cells.

LDL particles can associate with lectins, such as the jack bean lectin ConA, and this interaction can strongly reduce the proteolytic degradation of the lipoprotein particles in cells [48, 49]. When incubating human fibroblasts with LDL containing CTL-esters in the presence of ConA for either 3 or 6h, we found a large increase in PM fluorescence of the sterol at the expense of a reduced uptake of the LDL particles (Figure S3). Notably, the PM fluorescence recorded in the channel for CTL was heterogeneous and patchy, almost to the extent found in cells loaded with LDL containing CTL-ether (compare Figure S2C with the inset in Figure S3A).

These results show that the sterol fluorescence in the PM of non-treated cells consists of contributions from bound LDL particles containing CTL-ester as well as of hydrolyzed CTL exported from LE/LYSs to the PM. While our current setup does not allow us to distinguish these populations unequivocally, we surmise that a homogeneous PM staining stems from membrane-intercalated CTL, while LDL-particles bound to their receptor on the cell surface result in a patchy signal of CTL-esters in the LDL core. Supporting that notion, we have compared the membrane fluorescence of fibroblasts incubated with either Alexa488-LDL or LDL containing CTL-ester to that of cells stained with CTL delivered from albumin. We confirmed that the PM signal of Alexa488-LDL and of CTL derived from LDL is patchy after 2h of incubation, while that of CTL is homogeneous in the PM after overnight labeling from either LDL-associated esters or from albumin (Figure S4).

### Kinetics of lysosomal transport of LDL and of metabolism of its CTL-esters

Using quantitative live-cell fluorescence microscopy, we measured the kinetics of LDL transport to LE/LYSs containing Rh-dextran. We found that the sterol fluorescence from LDL containing CTL-esters increases in endo-lysosomes in a slightly sigmoidal fashion over time (Figure 3A and B). Such sigmoidal kinetics is well described by a sequential transport model (Figure 3B, inset and Materials and methods). The rate constants for the first and second phase (i.e., *k*_1_ and *k*_2_) were first determined by standard non-linear regression of this sequential model to the lysosomal accumulation of CTL/CTL-esters. We found the two rate constants to be equal and corresponding to a half time of 72.3 min for each step (see Tab. S3). To ensure that the measured CTL fluorescence in endo-lysosomes indeed comes from LDL-derived CTL-esters, we labeled the LDL particles additionally with an Alexa488-fluorophore on the apoB portion after reconstituting the CTL-esters and incubated cells with such double-labeled LDL particles for 24h (see Materials and methods). Unfortunately, the second labeling step strongly reduced the observable fluorescence of the sterol in cells, which made that this experiment is limited to the long incubation period of 24h to enable sufficient cellular uptake. Still, we can show that the remaining fluorescence derived from CTL-esters coincides with the Alexa488-fluorescence in intracellular vesicles located in the perinuclear area (Figure 3C). These results support that the majority of the lysosomal fluorescence of the CTL(-esters) at early time points is associated with LDL particles.

**Figure 3.**
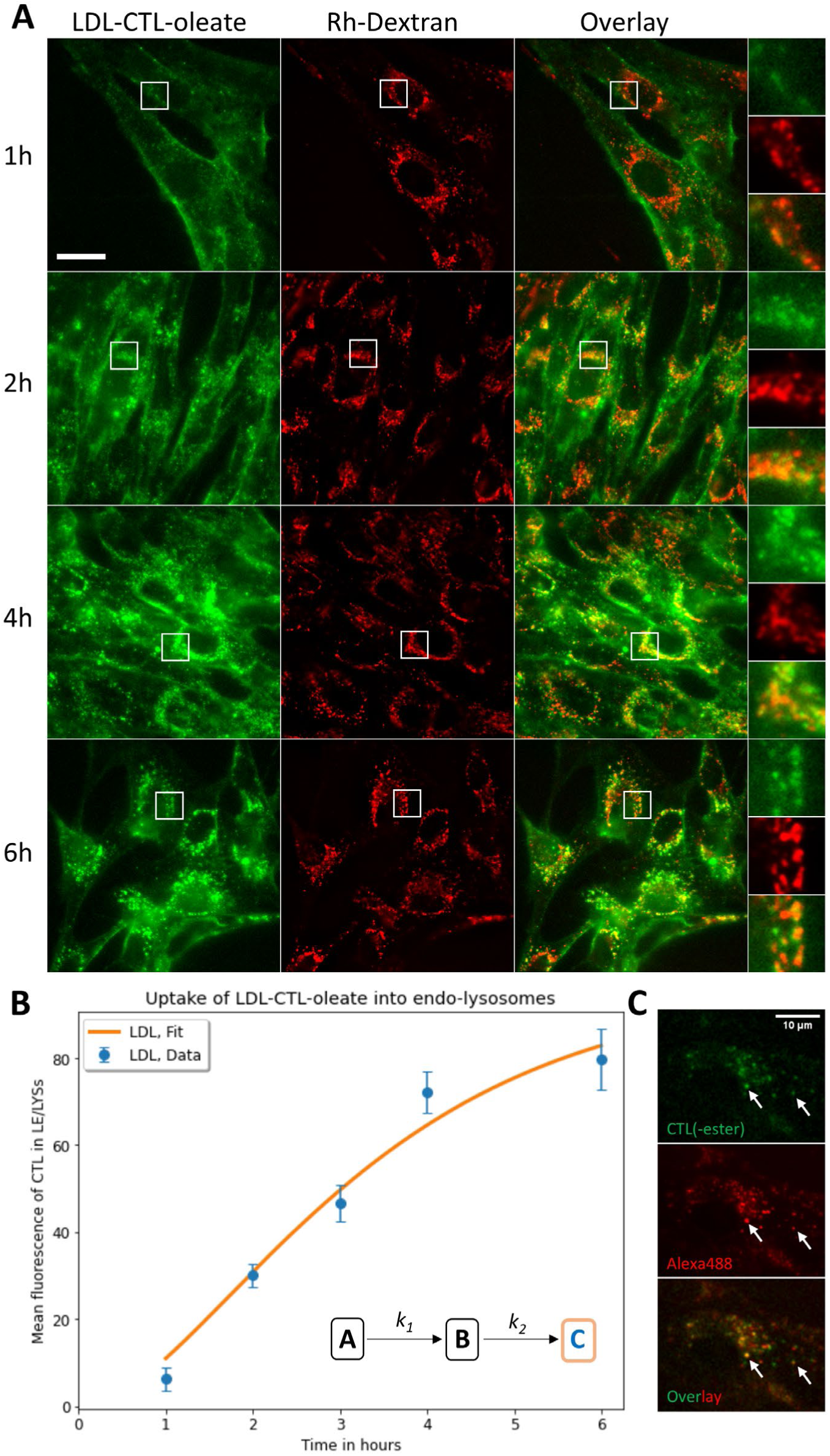
Kinetics of lysosomal accumulation of LDL-derived CTL(ester). Fibroblasts were incubated in LPDS medium supplemented with 0.5 mg/ml Rh-dextran overnight. On the following day, the cells were incubated in LPDS medium containing 0.1 mg/mL LDL with CTL-oleate ester for the indicated times. A, representative deconvolved images are shown for each time point with CTL ester fluorescence pseudocolored in green and Rh-Dextran in red. Co-localization appears orange. B, quantification of CTL ester fluorescence overlapping with the Rh-dextran signal in the endo-lysosomes and its kinetic modeling using a sequential transport model (see Materials and methods; blue dots, data mean +/- S.D., orange line, model fit) is shown. C, cells were incubated in LPDS medium with 0.2 mg/ml Alexa488-LDL containing CTL-oleate ester for 24h and imaged. Co-localization of LDLs apoprotein labeled with Alexa488 (green) with the fluorescent sterol, CTL(ester), pseudocolored in red, is indicated with arrows. Scale bar, 10 µm.

Using Lipid-MS, we were able to measure the kinetics of intracellular accumulation of hydrolyzed CTL derived from LDL-associated CTL-oleate esters as well as of re-esterification of the liberated CTL (Figure 4A and B).

**Figure 4.**
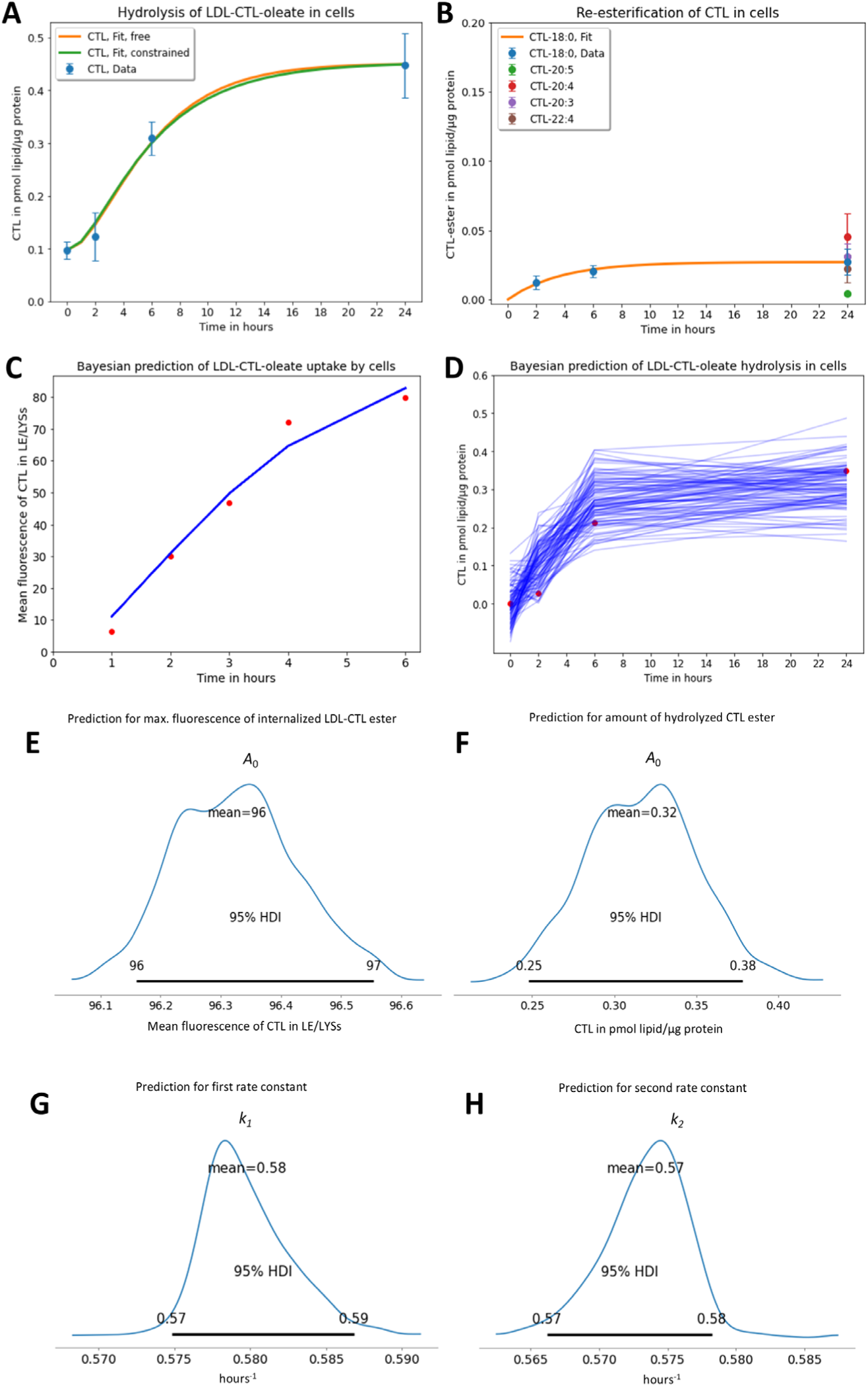
Lipidomic analysis of cells loaded with reconstituted LDL particles. Fibroblasts were incubated in LPDS medium containing 0.1 mg/mL LDL with CTL-oleate ester for the indicated times. A, hydrolysis of CTL-oleate ester was detected as a sigmoidal increase of the amount of free CTL over time (mean +/- S.D. of the data in blue). The data was fit to a sequential kinetic model, either with both rate constants, *k*_1_ and *k*_2_, allowed to vary (orange line) or with *k*_1_ being fixed at *k*_1_ = 0.0096 min^-1^ (green line). This is the value determined from the CTL(ester) transport data (compare Figure 3B and Tab. 1). B, re-esterification of CTL quantified for the indicated ester species (colored symbols at 24h). Only for the formation of CTL-stearate (18:0), a time course could be measured and is shown as mean +/- S.D. (blue symbols), together with a fit to a single exponential function (orange line, see also Tab. 1). C-H, hierarchical Bayesian modeling of the time courses for fluorescence increase (compare Figure 3B) and of the hydrolysis of LDL-associated CTL-ester in LE/LYSs. The mean values of the data are shown as red dots and the simulated time courses generated with parameters sampled from the posterior distribution as blue lines (n=100) for the fluorescence imaging data (C) and the Lipid-MS data (D), respectively. From the posterior distribution the values for the amplitudes (E and F) and shared rate constants *k*_1_ (D) and *k*_2_ (H) are shown with the 95% highest density interval (HDI) indicated.

**Table 1.** Kinetic parameters of CTL-ester transport and hydrolysis. Kinetic parameters are shown for the sequential model as determined by hierarchical Bayesian analysis (see Materials and methods). Amplitudes were determined separately, while the rate constant was shared for each data set. Values are given as mean ± standard deviation of the posterior distribution.

| Experimental data | Start amount in pool $A_0$ | Rate constant and half-time for first step | Rate constant and half-time for second step |
| --- | --- | --- | --- |
| Fluorescence intensity of LDL-derived CTL/CTL18:1 in LE/LYSs | $96.327 \pm 0.102$ a.u. | $0.0097 \pm 0.0011 \text{ min}^{-1}$<br>($t_{1/2} = 71.7 \text{ min}$) | $0.0096 \pm 0.0012 \text{ min}^{-1}$<br>($t_{1/2} = 72.6 \text{ min}$) |
| Concentration of LDL-derived CTL in LE/LYSs | $0.317 \pm 0.034$ pmol/$\mu\text{g}$ protein | | |

Formation of free CTL, indicating hydrolysis of the CTL-ester, was slightly sigmoidal, resembling that measured for lysosomal transport of these LDL particles (Figure 4A, compare with Figure 3B). To analyze this data further, we employed the same sequential transport model as used for quantifying the lysosomal fluorescence of CTL/CTL-esters. When fitting this sequential kinetic model to the hydrolysis kinetics, we found that the rate constant for the first step can be set to the value, we determined for the first step of the lysosomal accumulation of LDL, while the second step of the hydrolysis kinetics is slower with a half-time of 198.0 min (Figure 4A data, blue symbols; green line is fit to this model). These values were obtained using a non-linear regression routine, which only gives point estimates for the parameters and asymptotic confidence intervals. To better account for the variability and sparsity of the measured data, a Bayesian model approach is more suitable, as it provides information about model uncertainty and is more flexible when analyzing multiple datasets with shared parameters (see below) [50]. We therefore used the kinetic two-step model in a Bayesian framework (see Materials and methods for details). Since the data we collect is inherently noisy, we assume that the intensities we measure are part of a normal distribution with a mean equal to the average intensity of the determined kinetics and an unknown standard deviation, σ. Bayesian modeling of the sequential model provides not only the mean but the entire posterior distribution of the estimated kinetic parameters. This posterior distribution is the conditional probability of the parameter values given the observed data. It has as many dimensions as there are estimated parameters and allows for simulating time courses by sampling from it by Markov Chain Monte Carlo (MCMC) simulations (Figure 4C-H and S5). Using the Bayesian approach for model parameter inference on the fluorescence kinetics data of LDL transport to LE/LYSs, identical mean values as for the non-linear regression were found, and samples drawn from the posterior distribution are coinciding well and in a narrow range with the experimental data (Figure S5A-E and Tab. S3). By applying the same model separately to the time course of hydrolysis of LDL-derived CTL-ester, we found a very similar distribution for the first rate constant, but obtained a slower second kinetic rate constant with a broad distribution of values (Figure S5F-I and Tab. S3). The exact value of the second rate constant, *k*_2_, as inferred by Bayesian analysis is rather imprecise for the CTL-ester hydrolysis data, as the posterior is relatively wide with the 95% highest (posterior) density interval (95%HDI) varying from *k*_2_ = 0.2 to 0.62 s^-1^ (Figure S5I). Accordingly, the simulated time courses sampled from the posterior distribution by MCMC simulations spread much more around the measurement data than those obtained by Bayesian analysis of the CTL-ester uptake kinetics (compare Figure S5B and F).

The larger uncertainty for the second rate constant in the hydrolysis experiment is likely the result of the sparsity of the measurement data. This limitation can be overcome by using a hierarchical model in which the rate constants are shared between several data sets. We used this approach to determine the rate constants of the kinetic two-step model using the fluorescence kinetics of endo-lysosomal accumulation of LDL-derived CTL (Figure 3B) as well as the hydrolysis kinetics of LDL-derived CTL-esters (Figure 4A, symbols) as input. With such a hierarchical Bayesian model, we could increase the confidence in the parameter estimates and confine the time courses sampled from the posterior closer to the measured data (see Materials and methods for details and Figure 4C-F). This Bayesian analysis confirms the parameter values obtained by non-linear regression and additionally reveals that the variance of these parameter estimates is relatively low, suggesting that the model describes the data well (Tab. 1 and Figure 4C-F).

What are the cellular processes underlying these two kinetic steps of LDL processing? Binding of LDL to the cell surface, its transport to sorting endosomes (SEs), as well as dissociation of LDL from its receptor inside early endosomes are fast events with half-times of 2-5 min [10, 11, 15, 51]. Thus, the measured kinetics of accumulation of LDL containing CTL- ester in endo-lysosomes is likely not limited by these processes. We hypothesize that the initial arrival of LDL in late endosomes resembles the first kinetic step, which has previously been shown to be the consequence of slow maturation of early into late endosomes [10, 11, 52, 53]. The second step is likely a combination of continuous transport of the LDL containing CTL- oleate into LE/LYSs, eventually also via other pathways than that mediated by the LDL-R [47], and of degradation of the LDL particles inside these organelles. The latter is supported by our observation that CTL-esters are partially quenched in the LDL particles, as addition of the detergent Triton X100 to reconstituted particles resulted in an increase of the CTL signal (Figure S1C). The overall half-time of the sequential hierarchical Bayesian model with shared kinetic parameters is 3h for the accumulation of CTL fluorescence in LE/LYSs and for the hydrolysis of the CTL-oleate (Figure S6). These results can be compared to values in the literature, which used LDL containing radio-labeled CEs and cell fractionation to show that hydrolysis of CEs takes place after an initial lag-phase. From such studies, the half-time of hydrolysis can be estimated by inspection of the respective graphs with estimated values ranging from about 1.5-2h in CHO cells [17, 18, 54] and 2-6h in primary human fibroblasts [14, 55, 56]. The latter are also in agreement with the sequential transport kinetics described for LDL-R dependent and independent transport of radiolabeled LDL tagged at the apoprotein in primary human fibroblasts [15, 46, 47, 55]. Thus, a sequential transport model with the first step describing LDL uptake and the second step CTL-ester hydrolysis/dequenching is a good description for the trafficking of LDL-derived CTL esters in human fibroblasts. The model predicts that degradation of LDL’s CEs first takes place once the particles have reached LE/LYSs. Only in these organelles, the pH is sufficiently acidic for acid lipases to be active in hydrolyzing LDL-derived sterol esters. This conclusion is supported by the experiments with Lalistat (see above).

Lipid-MS also allows us to quantify the re-esterification of CTL, but only as long as we measure formation of other esters than CTL-oleate. That is, re-esterification to oleate cannot be distinguished from CTL-oleate taken up by cells as part of LDL, so re-esterification must be assessed by formation of other CTL-esters in the cells. Indeed, we found that cells form CTL-stearate (i.e., CTL-18:0) with mono-exponential kinetics and a half-time of 152.7 min (Figure 4B and Tab. S3). After prolonged incubation, also CTL-esters with polyunsaturated fatty acids, such as CTL-20:4, could be measured (Figure 4B, red symbol). These results show that hydrolysis of CTL-oleate and re-esterification of the freed CTL take place on comparable time scales, suggesting that transport of the liberated CTL to the ER, where ACAT resides, is rapid.

### LDL-derived cholesteryl esters and triacylglycerols can be tracked by lipidomics

Fibroblasts accumulated primarily sterols during uptake of LDL, as expected given the lipid composition of the LDL particles (Figure S1 and S7, red symbols). This allows us to follow the dynamics of specific CEs, which are enriched in LDL particles but not abundant in cells. In particular, we found that the species CE16:0, CE16:1, CE18:1 and CE18:2 are very abundant in LDL but not in cells, enabling us to trace these CEs specifically in cells (Figure S8). The species CE18:3 is isomeric to CTL18:1 and was therefore not included in this analysis. To analyze this data, a hierarchical mono-exponential Bayesian model was used, in which the amplitudes for all four species were independently determined, while the rate constant was shared for all four time courses. The inferred rate constant was *k* = 0.00358 ± 0.00003 min^-1^, corresponding to a mean half-time for cellular accumulation of all studied LDL-specific CEs of *t*_1/2_ = 193.44 min. A mono-exponential rise to maximal cellular accumulation with an exponential increase and comparable half-time was also found in previous work using radioactive CE tracers in LDL [14]. Since the CEs, we determined, are likely a mixture of LDL-derived internalized CEs and hydrolyzed but reesterified cholesterol, we cannot further distinguish individual pools in this experiment, in contrast to the studies with the reconstituted CTL18:1 ester (see above).

In addition to CEs, we found that LDL also contains specific TAGs, which are of very low abundance in the non-labeled cells (Figure S9). These TAGs can also be used to track the uptake of LDL-derived core lipids. Using the hierarchical monoexponential Bayesian model with shared rate constant, we found that a mono-exponential rise to the maximal value also describes the cellular accumulation of LDL-specific TAGs very well, just as we observed for the LDL-associated CEs (compare Figure S8 and S9). The inferred rate constant was with *k* = 0.00168 ± 0.00008 min^-1^ lower than that of the accumulation of CEs and corresponds to a half-time of *t*_1/2_ = 411.8 min. The slower time course of TAG enrichment in cells compared to that of CEs could be the result of more extensive remodeling, with part of the TAG-derived fatty acids used in β-oxidation and phospholipid synthesis. Together, our results show that lipidomics combined with Bayesian kinetic modeling is very powerful in tracing different lipoprotein-derived lipid species in cells.

### Remodeling of glycerophospho- and sphingolipids during uptake and processing of LDL

Apart from these changes, we observed alterations in the abundance of phospho- and sphingolipids. Since these lipids are less specific in LDL compared to cells, we could not trace the accumulation of particular species unequivocally but rather assess an overall change in their abundance upon incubating cells with LDL particles. We found a slight decrease in glycerophospholipids due to the initial 6h of incubation, which was followed by an increase after 24h (Figure S7, green symbols). *De novo* synthesis of glycerophospholipids to provide membrane lipids for cell division likely contributes to the long-term increase of glycerophospholipids. This was also found for various species of PC (Tab. S1 and Figure S10A). A cellular response to the uptake of LDL is a drop in lysoPC, probably due to the reacylation (Figure S10B). While the overall abundance of SM increased only slightly, there was a strong formation of ceramides, particularly at later time points (Figure S11). LDL particles contained mainly SM 34:1,2, SM 40:1,2 and SM 42:2,2, and we therefore tracked those SM species over time in cells (Tab. S2 and Figure S11, dashed lines). The corresponding ceramide species increased over time (Figure S11, straight lines of the same color). This implies the activity of acid sphingomyelinase (e.g., SMPD3), degrading the LDL-associated SM inside LE/LYSs [57, 58]. The cellular abundance of PE species followed that of PC, with species-dependent remodelling of lyso-PE (Figure S12A and B). Cell-associated diacyl-PE ethers also showed an initial drop followed by a slight increase for most species, likely due to de novo synthesis in connection with cell division (Figure S12C). Together, these results show the power of lipidomics to uncover cellular accumulation and metabolic conversion of lipoprotein-associated lipids. Together with the live-cell imaging of LDL-associated CTL- esters, these results demonstrates that Lipid-MS also provides detailed insight into remodelling of the cellular lipidome upon lipoprotein uptake.

### Endo-lysosomes containing LDL align with the endoplasmic reticulum for sterol transfer

Given that we observed re-esterification of LDL-derived CTL already after 6h of incubation, we hypothesized that hydrolysis of the CTL-esters inside LE/LYSs is followed by the immediate export of the generated CTL to the ER. Since the mole fraction of cholesterol in the ER is low, and the membrane area of this organelle is very large, direct visualization of CTL in the ER is challenging. Nevertheless, already after 6h of loading cells with LDL containing CTL-ester, we found that LE/LYSs containing the fluorescent sterol and the organelle marker Rh-dextran overlap with ER-tubules labeled with NBD-ceramide (Figure 5A). NBD-ceramide is a well-known marker for the trans-Golgi network, but it also gives a faint staining in the ER, since it is transported between both compartments [59, 60]. We can confirm this by confocal imaging of fibroblasts double-labeled with NBD-ceramide and ER-tracker (Figure S13). Neither after 6h nor after 24h of incubation, CTL was transported to the TGN, which was easily identified by its bright staining with NBD-ceramide next to the nucleus (Figure 5A, B, yellow arrows). After incubating fibroblasts for 24h with LDL containing CTL-esters, we found some sterol fluorescence in lipid droplets stained with Nile Red (Figure 5C). Formation of droplets in LDL-loaded human skin fibroblasts has also been found previously [61], and droplet-associated CTL fluorescence is in line with our results of the Lipid-MS, demonstrating that the hydrolyzed CTL become re-esterified and stored in droplets. After 6h of incubation with LDL containing CTL-esters, most sterol fluorescence in the PM is homogeneous, suggesting that it stems from hydrolyzed CTL exported to the cell surface.

**Figure 5.**
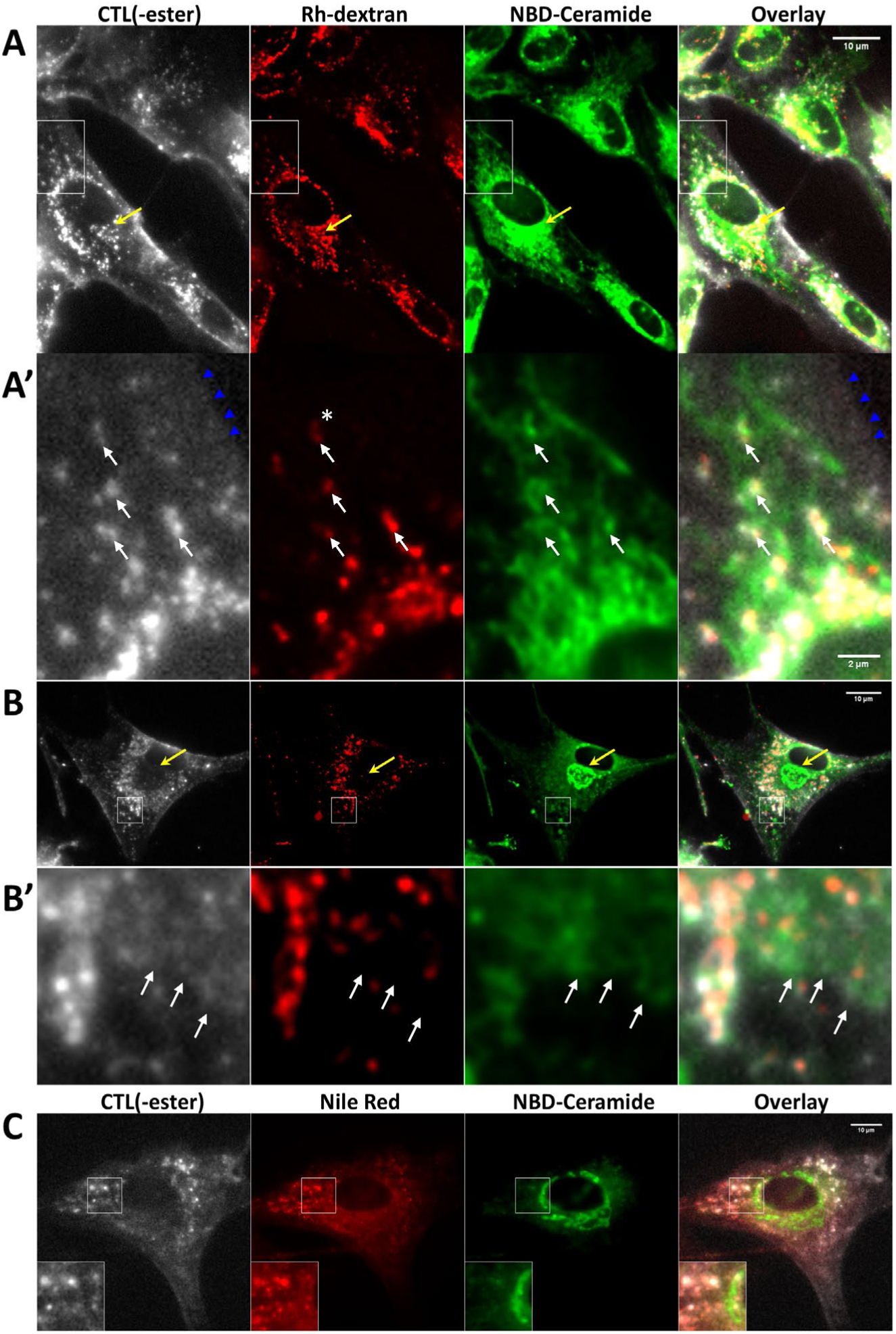
Fluorescent sterols released from LDL particles co-localize with the ER and LDs. Fibroblasts were incubated in LPDS medium containing 0.1 mg/mL LDL with CTL-oleate ester for either 6h or 24h. LE/LYSs were stained with 0.5 mg/ml Rh-dextran overnight, and the ER and Golgi apparatus were labeled with NBD-ceramide added during the last 10 min of incubating cells before imaging. After 6h (A and A’), endolysosomes containing CTL(ester) and Rh-dextran were aligned with ER tubules which contained NBD-ceramide (white arrows). Faint CTL fluorescence in LE/LYSs overlapped occasionally with the ER (white star, only shown in Rh-dextran panel in A’). Sterol fluorescence in the PM is indicated with blue arrowheads. After 24h of incubation (B and B’), faint fluorescence of CTL(ester) was additionally found co-localizing with the NBD-ceramide positive ER structures (white arrows). C, in addition to the co-staining with NBD-ceramide, the droplet marker Nile Red was added during the last 10 min of the 24-h incubation period. CTL(ester) fluorescence was found in LDs (inset in C). Neither after 6h of incubation nor after 24h, CTL(ester) fluorescence was found in the Golgi apparatus, localized next to the nucleus (yellow arrows in A and B).

We found in independent experiments using a spinning disk confocal microscope that endo-lysosomes containing the Cathepsin B substrate MagicRed form frequent contacts to the ER, in line with earlier reports (Figure 6A for examples) [62]. We chose MagicRed for these experiments, as its strong red fluorescence is only found in hydrolytically active endo- lysosomes [63]. Using our recently developed automated image analysis protocols (see Materials and methods and [64]), the distance between those endo-lysosomes and the ER was quantified from 3D confocal stacks. For comparison, the organelle size was measured and determined as the average radius of LE/LYSs (i.e., the distance from the 3D centroid position to the endo-lysosome surface). This distribution peaks around 0.3 µm with a long tail up to about 1.5 µm and a slight shift to larger sizes in LDL-treated cells (Figure 6B). The closest distance of the centroid of endo-lysosomes to ER tubules was about 0.7 µm in the absence of LDL but about 0.3 µm in the presence of LDL (Figure 6B and see Fig. S14 for the corresponding cumulative frequency plots). Since the latter matches the organelle size, we can conclude that endo-lysosomes form contacts with the ER upon ingestion of LDL.

**Figure 6.**
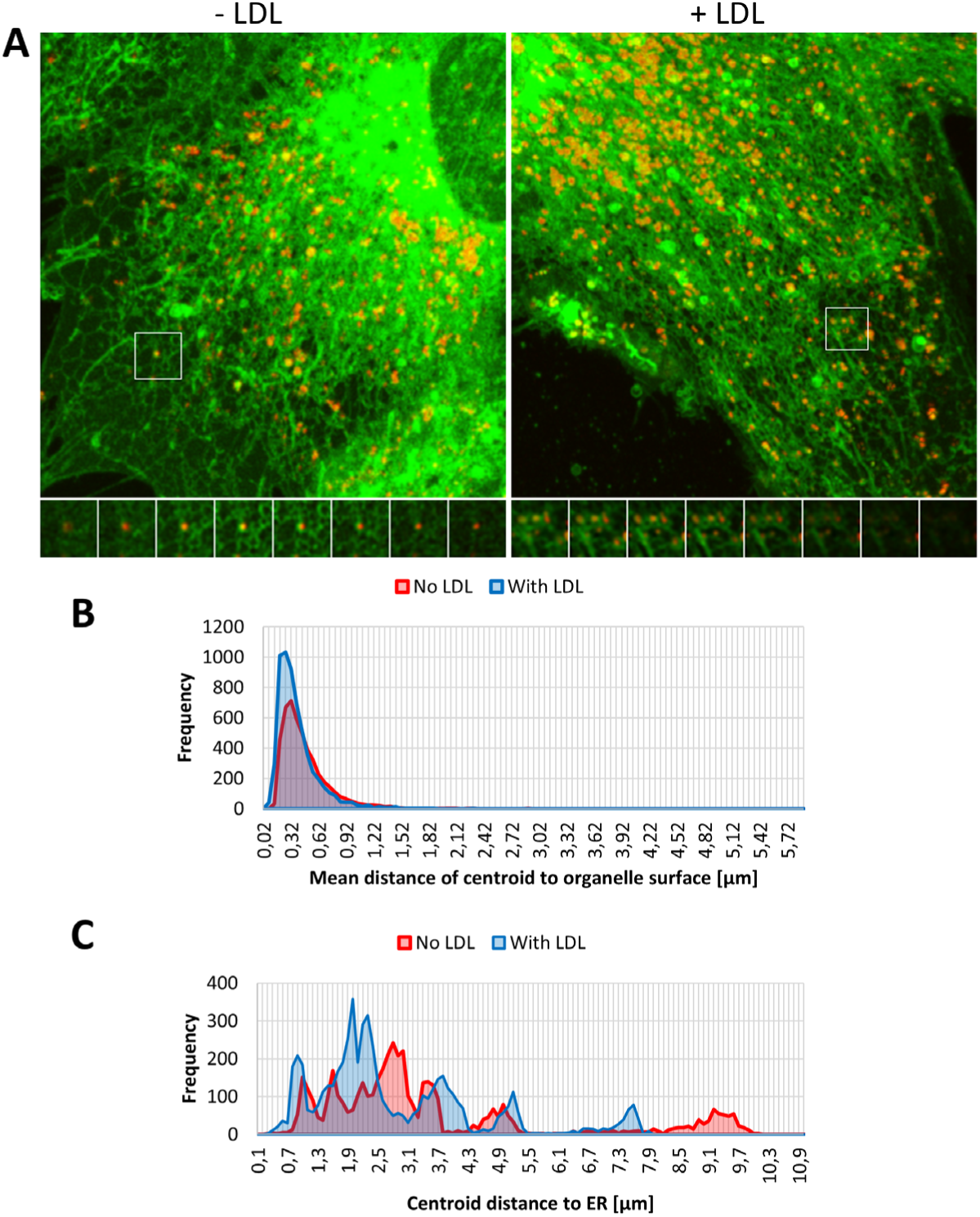
Internalized LDL mediates recruitment of endo-lysosomes to the ER. Fibroblasts were incubated in LPDS medium containing 0.5 mg/ml MagicRed (red) overnight followed by incubation with 0.1 mg/mL LDL (right panel in A, ‘+ LDL’) or without LDL (left panel in A, ‘-LDL’) for 6h. During the last 10 min of incubation 5 µM NBD-Ceramide were added to label the ER (green). 3D image stacks were acquired at a spinning disk confocal microscope, deconvolved and further analyzed to determine the distance between MagicRed stained LE/LYSs and NBD-Ceramide-labeled ER tubules (see Materials and methods). B, C, histogram of endo-lysosome size (B) and distance to the ER (C) for cells incubated without LDL (red) or with LDL (blue). The mean distance between the particle centroid and the organelle surface is a measure of endo-lysosome size, while the shortest distance between endo-lysosome centroid and the next ER tubule quantifies ER-LE/LYSs distance. For this analysis, at least eleven 3D stacks of 1-2 cells with 350-620 endo-lysosomes in each stack were quantified for each condition.

Moreover, LDL tagged with a fluorescent dye on the apoprotein moiety (Alexa488-LDL) was often found in close proximity to the ER (Figure 7A): single particle tracking demonstrates that Alexa488-LDL containing endo-lysosomes stayed in contact with ER tubules for several seconds per event, where their mobility is reduced compared to freely moving vesicles (Figure 7B-J and Supplemental video 1). From the videos, the distance between the surface of LE/LYSs containing Alexa488-LDL and the skeletonized ER tubules was measured. This distance was often found to be between 0.05 and 0.2 µm, indicating contact formation between both organelles. Together, these results demonstrate that LE/LYSs become recruited to the ER upon ingestion of LDL, and they suggest that membrane contact sites between endo-lysosomes and the ER could facilitate transfer of LDL-derived cholesterol to the ER for esterification.

**Figure 7.**
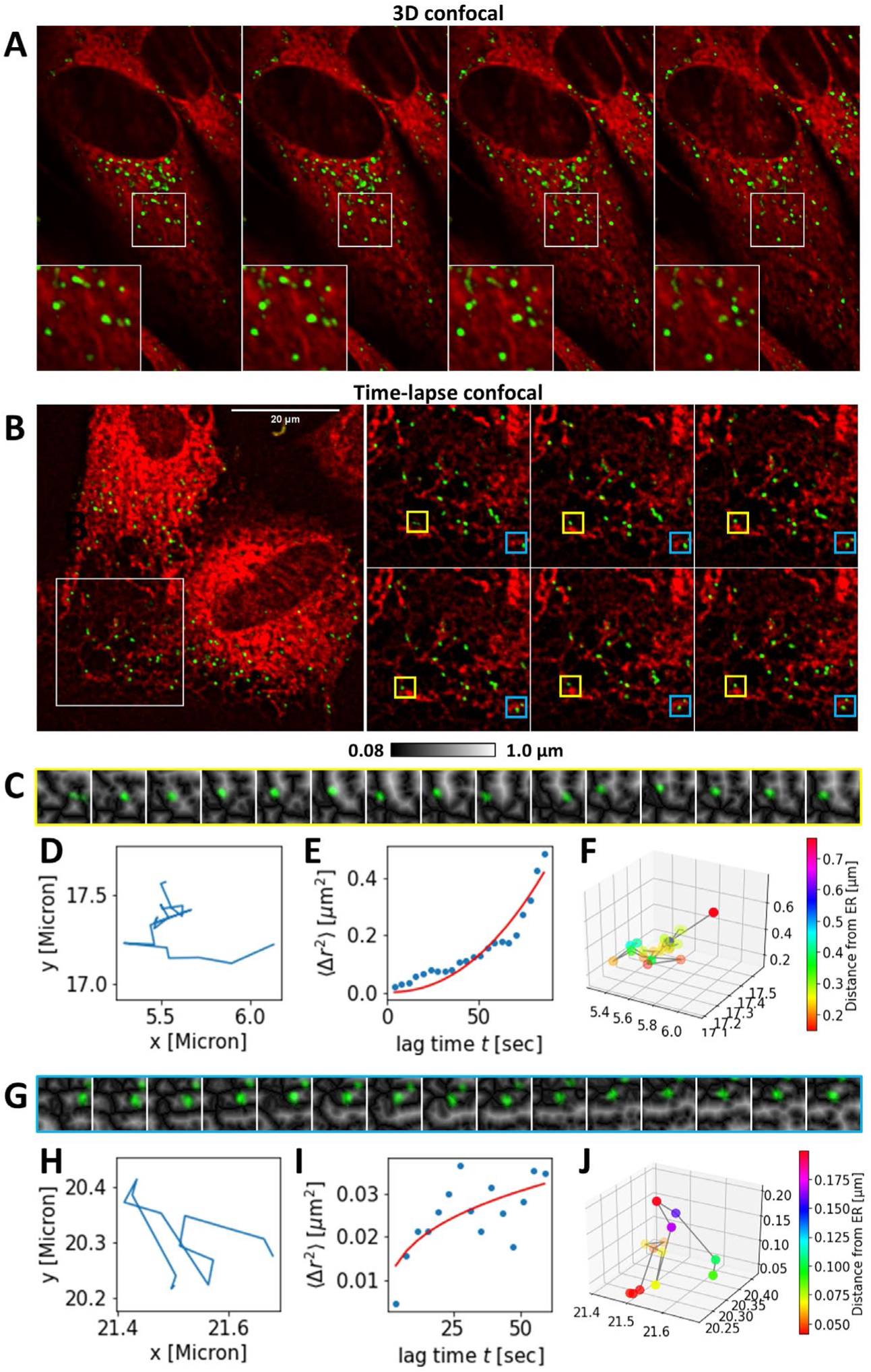
Endolysosomes containing fluorescent LDL form frequent contacts to the ER. Fibroblasts were incubated in LPDS medium overnight, followed by washing with PBS, incubation with 0.5 mg/ml Alexa488-LDL for 1.5h and staining with ER tracker for the last 30 min of incubation. Cells were washed and imaged on a spinning disk confocal microscope, deconvolved and further analyzed to determine the distance between LE/LYSs containing Alexa488-LDL (green) and ER Tracker Red-labeled ER tubules (see Materials and methods). A, montage of 3D stack acquired along the optical axis every 0.25 µm with Alexa488-LDL in green and ER Tracker Red in red. B-J, time-lapse imaging for the cells acquired every 3.9 sec. B, an overview image with selected frames of the rectangular box in the left panel. Two regions of interest were selected (yellow and blue box) of which the Euclidian distance map was calculated, as shown in C and G. LE/LYSs containing Alexa488-LDL were tracked by single particle tracking, and their trajectories (D and H), their mean squared displacement (MSD; E and I, blue dots) and their distance to the ER (F and J) were calculated. The MSD was fit to an anomalous diffusion model (E and I, red line). See the main text and Supplemental materials for further details.

## CONCLUSION

In this study, we present a novel approach to study the trafficking and metabolism of LDL-derived cholesterol in living cells. Our new method is based on intrinsically fluorescent sterol esters, which differ from natural CEs only by having two additional double bonds in the steroid ring system. This minimal modification ensures close resemblance to the natural cholesteryl oleate. We also synthesized a CTL-ether, which cannot be hydrolyzed by lipases, as shown *in vitro* and in cells. Binding of both LDL containing CTL-ester and -ether could be outcompeted by excess unlabeled LDL to a similar extent as previously described for ^125^I-labeled LDL in human fibroblasts [15]. In line with these earlier studies, we conclude that several binding sites contribute to the uptake of LDL, with the LDL-receptor representing the major one. Using Lipid-MS, we demonstrate that LDL-derived CTL-esters become hydrolyzed in endo- lysosomes of human fibroblasts after an initial delay phase. In comparison, re-esterification of some of the CTL in the ER, as well as CTL transport to the PM, are fast events, suggesting that trafficking of LDL to LE/LYSs and lysosomal hydrolysis of LDL-associated CEs are the rate-limiting steps in redistributing LDL-derived cholesterol in human fibroblasts. We have developed a hierachical Bayesian framework, which allows us to infer rate constants for a sequential model, which accounts for both processes and includes uncertainty quantification using imaging and lipidomics data as model input. We show the strength of this to our knowlegde novel data analysis approach by also quantifying the uptake kinetics of various LDL-derived CEs and TAGs. Accumulation of LDL-derived CEs follows mono-exponential kinetics with a mean half time of a little more than 3h. Since we observed re-esterification of CTL derived from hydrolyzed LDL-CTL esters with a half-time of just 2.5h, we conclude that cellular accumulation of CEs is accompanied extensive acyl chain remodeling. Re-esterification of CTL was paralleled by LDL-induced, apposition of endo-lysosomes with the ER, suggesting that transport of cholesterol derived from hydrolysis of CEs to the ACAT-site in the ER takes place via membrane contact sites forming between LE/LYSs and the ER, as suggested earlier [21, 22, 65–67]. Tethering of endo-lysosomes to the ER is a tightly regulated process, including sorting nexins, such as SNX13 and SNX19, and the sterol transporters StARD3 and Aster B (GRAM1B) [40, 68–70]. In a recent study, NPC1 has been shown to regulate contacts with the ER to mediate lysosomal cholesterol egress in concert with the ER-resident GRAM1B [70]. Similarly, the oxysterol-binding protein ORP1L has been associated with tether formation, endo-lysosome positioning, and trafficking of cholesterol from LE/LYSs to the ER [34–36, 70]. It is likely that one or several of these transporters get activated by ingested LDL, which could explain our findings of the recruitment of LE/LYSs to the ER upon LDL loading. On the other hand, SNX13 has been shown to be a negative regulator of NPC1- mediated cholesterol export, and its absence likely redirects cholesterol transport to the PM independent of NPC1 [69].

Other ER-resident proteins, such as the Aster/GRAMD1 family, have been implicated in the export of LDL-derived cholesterol to the PM [24, 43]. Our current setup does not allow us to quantitatively distinguish between the CTL signal derived from CTL-esters as part of LDL particles bound to the cell surface and the fluorescence of CTL transported to the PM upon lysosomal hydrolysis of the esters. This is particularly relevant at early time points up to 6h, when a large portion of LDL still resides at the PM before equilibrating with intracellular pools, while lysosomal hydrolysis of CTL-oleate and, as a result, also export of CTL from LE/LYSs has started [16]. Surface-bound LDL particles containing CTL-oleate esters give a patchy CTL signal in the PM, which resembles that of LDL particles. After 24h of incubation with LDL containing CTL-esters but not with LDL containing CTL-ethers, we found predominantly a homogeneous membrane signal of CTL in control cells, indicating that this originates from free CTL exported to the PM. Inhibiting acid lipase with Lalistat abolished this homogeneous membrane staining, indicating that LAL is the major enzyme responsible for the hydrolysis of LDL-derived CEs in fibroblasts, in agreement with other reports [14]. Re-esterification of hydrolyzed CTL, however, follows immediately the lysosomal degradation of the CTL-esters, such that it is possible that LDL-derived cholesterol destined for the PM traffics via the ER as a transit compartment. Since cholesterol has a much higher affinity for saturated phospholipid species found in the PM compared to the more unsaturated species of the ER, the PM could act as a sink for the lipoprotein-associated cholesterol, even when it passes through the ER network [2, 71]. Future studies are warranted to shed light onto pathways for trafficking of LDL-derived cholesterol to the cell surface.

## MATERIALS AND METHODS

### Reagents

The human lipoprotein depleted serum (LPDS, S5519), LDL (L8292), cholesteryl oleate (303- 43-5), Sigmacote (SL2), potato starch (9005-25-8), U18666A (U3633), Lalistat-2 (SML2053) and concanamycin A (ConA) (27689) were purchased from Sigma Chemicals (St. Louis, MO). Fetal bovine serum (FBS) and DMEM were from GIBCO BRL (Life Technologies, Paisley, Scotland). Rhodamine-labeled dextran (D1819), ER-tracker red (E34250), Nile Red (N1142), Alexa Fluor 488 Protein Kit (A10235) and NBD-C6-Ceramide (N1154) were from Invitrogen/Molecular Probes (Inc. USA). The Amplex Red kit (A12216) for quantification of cholesterol was purchased from ThermoFisher (Denmark office, Odense, DK). CTL, CTL- oleate ester and ether were synthesized as described in Supplemental materials.

### In-vitro hydrolysis experiments of fluorescent cholesteryl esters

The Amplex Red kit for quantification of cholesterol contains a cholesteryl esterase, which was used to assess the hydrolysis of CTL-oleate ester relative to its natural counterpart, cholesteryl oleate. First, standard curves of either cholesterol or CTL were generated using cholesterol oxidase-induced production of hydrogen peroxide. The latter is detected by horsradish- peroxidase induced formation of resorufin from the Amplex Red reagent, following the manufacturer’s instructions. Since we found that the oxidation efficiency of the CTL was less efficient than that of cholesterol, two separate standard curves were required. Next, 2.5 or 5 µM of either CTL 18:1 ester or CTL 18:1 ether were dissolved in the reaction buffer of the Amplex Red kit, and the reaction was initiated by adding the reaction mix containing the esterase, followed by an incubation period of 30 min at 37°C. Finally, the fluorescence intensity was quantified using a microplate reader in an excitation range of 530-560 nm and an emission wavelength of 590 nm. Since the emission of CTL 18:1 ester and ether is in the UV region of the spectrum, no interference with the AmplexRed fluorescence readout was found. In the second assay, the sterol esters and ethers were incubated with the esterase from the Amplex Red kit for the recommended amount of esterase, and the reaction mixture was incubated for 2h at 37°C. Subsequently, the mixtures were spun down in the vial at 150 rpm, and lipids were extracted with a 10-fold excess of chloroform/methanol (3:1) while vortexing the solution for 2 min. After centrifugation at 2000 g for 4 min at room temperature, the lower phase was collected, evaporated, and resolubilized in 50 µL chloroform before spotting an aliquot onto a silica TLC plate alongside reference lipids. Petroleum ether: ethyl acetate (4:1, v/v) was used as the running solvent. To identify the sterols, the plates were treated with copper sulfate, as described [72]. Exposure to the heat emitted by a hot air dryer revealed the lipids as distinct brown bands. Fluorescence imaging of the plate was performed at 302 nm to confirm the presence of CTL incorporation (not shown).

### Cell culture

Human skin fibroblasts from a healthy male donor (Coriell Institute #GM08680) were cultured in T25 culture flasks, at 37 °C in an atmosphere of 5 % CO2 in a complete DMEM culture medium supplemented with 1 % glutamax, 1 % Penicillin-Streptomycin, and 10 % FBS. Cells were checked daily and split with trypsin when a confluency of 90 % was reached. Before fluorescent microscopy, cells were placed on 35 mm microscope dishes with glass bottoms (P35G-1.5-50-C, MatTek) and allowed to settle for 24-48 h in their culture medium before initiating experiments.

### Reconstitution of LDL with fluorescence sterol ester or ether

LDL reconstitution with CTL 18:1 ester was performed based on the method developed by Krieger et al. (1978) with minor modifications [73]. Briefly, LDL (2 mg; SAE0053, Sigma) was combined with potato starch (25 mg; S4251, Sigma) in Sigmacoated glass vials (SL2, Sigma). The mixture was snap-frozen in liquid nitrogen, held at -80°C for 2 hours, and lyophilized overnight at -110°C. Endogenous LDL lipid core extraction involved three rounds of vortexing 5 mL ice-cold heptane with lyophilized LDL, centrifuging at 5000×g for 10 min at 4°C, and removing the supernatant. Post-extraction, LDL was reconstituted with 1 mg CTL oleate ester and 5 mg cholesteryl oleate dissolved in 200 μL ice-cold heptane, gently mixed, and incubated for 10 min at -20°C. Heptane was evaporated under nitrogen flow, and particles were solubilized overnight in 10 mM Tricine buffer (pH 8.4) at 4°C.

Reconstituted LDL was isolated by sequential centrifugation: 5000×g for 10 min (glass vial), 9000×g for 10 min (Eppendorf tube), and repeated twice with the supernatant transferred each time. The resulting LDL particles containing either CTL-oleate ester (LDL-CTL-ester) or CTL- oleate ether (LDL-CTL-ether) were further characterized by measuring their emission spectra (Figure 1B). For that, the labeled particles were diluted in PBS, and fluorescence spectra were recorded using an ISS K2 spectrofluorometer (ISS, Champaign, IL, USA) with a 300 W Xenon arc lamp (slit width: 0.5 mm). LDL-CTL was excited at 328 nm, and emission was measured from 350–500 nm. To confirm CTL oleate-ester/ether incorporation, lipids were extracted from the reconstituted LDL by mixing 1:10 with CHCl_3_: MeOH (2:1), resulting in a 2-phase system, of which the bottom layer was isolated and evaporated under a stream of N_2_. The resulting lipid film was resolubilized in 50 µL CHCl_3_ and spotted onto a silica TLC plate alongside reference lipids. Petroleum ether: ethyl acetate (4:1) was used as the running solvent. Fluorescence imaging of the plate was performed at 302 nm to confirm CTL incorporation.

### Preparation of LDL labeled with Alexa488 and LDL-CTL-oleate

Native LDL or LDL-CTL-oleate ester at a concentration of 2 mg/ml were labelled with Alexa488-TFP-ester mixed with 50 µl of 1 M sodium bicarbonate buffer, following the manufacturer’s instructions. To remove unbound dye, the solution was dialyzed with PBS overnight in a dark, cold room maintained at 4°C.

### Co-labeling of cells for trafficking studies

Cells were prepared as described above, and before imaging, the culture medium was changed to DMEM containing LPDS, in which cells were incubated for 24 h. Subsequently, cells were labeled with the desired fluorescent version of LDL for the given period of time (indicated in the Results section). Prior the microscopy, cells were loaded with either 0.5 mg/mL Rh-Dextran for 24 h or with MagicRed to stain LE/LYSs. While Rh-dextran is a content marker of endo- lysosomes, MagicRed is a substrate for Cathepsin B, which fluoresces upon cleavage, indicating hydrolytically active LE/LYSs [63]. For co-staining the ER and Golgi apparatus, cells were incubated with C6-NBD-Cer for 10 min at 37°C (final concentration was 5µM). In some experiments, the ER-tracker (final concentration 1 µM) was loaded for 30 min prior to imaging (see more information in the Results). Lalistat was added to the microscopy dish (final concentration 10µM) 6h before loading with the fluorescent LDL, and the inhibitor was present during the subsequent overnight incubation.

### LDL vs LDL-CTL competition assay

Primary fibroblasts grown on microscope dishes were incubated in LPDS medium overnight. The following day, 1 mL LPDS medium was mixed with 0.5 mg/mL Rh-dex, 0.05 mg LDL- CTL-ester or ether, and with or without 4 mg unlabeled LDL. This mixture was given to the cells and incubated for 6 h at 37° C, 5 % CO_2_. The cells were washed with PBS before they were imaged in M1 buffer medium (150 mM NaCl, 5 mM KCl, 1 mM CaCl_2_, 1 mM MgCl_2_, 5 mM glucose and 20 mM HEPES (pH 7.4)).

### Fluorescence microscopy

A Leica DMIRB microscope with a 63x 1.4 oil immersion objective (Leica LAsertechnik GmbH) controlled by a lambda SC smart shutter (Shutter Instrument Company) was used to carry out widefield fluorescence microscopy. The microscope was equipped with an Andor IxonEM blue EMCCD camera operated at -75° C and driven by the Solis software supplied with the camera. CTL was imaged by the use of a specially designed filter cube from Chroma Technology (Brattleboro, VT, USA) with a 335-nm (20-nm bandpass) excitation filter, 365- nm dichromatic mirror, and 405-nm (40-nm bandpass) emission filter. The UV signal of CTL was always collected as bleaching stacks consisting of 50-100 frames. To correct for chromatic aberration, the focus plane was adjusted when acquiring images of CTL, as described previously [74]. The green fluorescence of Alexa488-LDL and C6-NBD-Cer was imaged with a standard fluorescein filter set with a 470-nm (20- nm bandpass) excitation filter, 510-nm dichromatic mirror, and 537-nm (23-nm) bandpass emission filter. For imaging the red fluorescence or Rh-dextran, Nile Red or ER-tracker, a standard rhodamine filter set with a 535- nm (50-nm bandpass) excitation filter, 565-nm dichromatic mirror, and 610-nm (75-nm bandpass) emission filter was used. Spinning disk confocal microscopy was carried out on a Nikon Ti-E spinning disc microscope with a 60×NA 1.4 oil objective, a Yokagawa CSU-X1 spinning disk and equipped with an Okolab microscope stage incubator to maintain the temperature at 37 °C and 5% CO2. The laser lines of 491 nm and 561 nm were used to excite C6-NBD-Cer, Alexa488-LDL and Er-tracker, respectively. An electron-multiplying CCD camera (Andor iXon EMCCD DU-885, 1004×1002 pixels, 8×8 μm) was used to acquire the images on that instrument.

### Image analysis

Deconvolution of images was done with the Richardson-Lucy algorithm implemented in ImageJ (http://rsb.info.nih.gov/ij) [185] with a theoretical point spread function calculated with the appropriate wavelength, refractive index, numerical aperture, and pixel spacing. Cellular autofluorescence in the UV channel was separated from the fluorescence CTL signal by subtracting the last frame from the first of the bleaching stacks [64]. The fractional fluorescence of CTL(ester) in LE/LYSs was calculated based on the binary image of the Rh-dextran images outlining the area covered by endo-lysosomes and normalized to the total cell associated intensity [75, 76]. Inter-organelle distances were measured by first calculating an Euclidean distance transform from the binarized ER image after appropriate thresholding, followed by localizing all endo-lysosomes in 3D stacks and determining their position on the Euclidean distance map, as described [64]. Tracking of membrane contacts between ER tubules and LE/LYSs containing Alexa488-LDL was carried out as described [64].

### Kinetic modeling and Bayesian model inference

Kinetic modeling of the data for transport and hydrolysis of LDL-associated CTL-esters was based on a sequential 2-pool model (see Supplementary notes for model equations), solved analytically and fitted to the experimental data using non-linear regression as implemented in the Scipy library in Python [77]. Alternatively, the data was fit to the same model in a Bayesian setting with normal distributions for the mean and a half-normal for the standard deviation as informed priors using the Python library PyMC3 [50]. The Bayesian model was defined either separately for both data sets or implemented as a hierarchical model with separate amplitudes but shared rate constants for the fluorescence-microscopy and Lipid-MS derived time courses. The posterior was calculated using Monte Carlo sampling for 1000 steps in one chain after tuning for 500 steps. Bayesian modeling of the cellular accumulation of CEs and TAGs was also based on a hierarchical approach but using a simpler mono-exponential 1-pool model (see Supplemental materials). Here, the amplitudes were inferred separately for each data set, while the rate constant was shared for all species of CEs and TAGs, respectively. MCMC simulations were carried out using the NUTS sampler with adaptive scaling for 1000 steps on 4 chains and an initial tuning phase for 500 steps. Random sampling from the posterior was used to simulate time courses for each condition.

### Lipidomics

Cells were prepared as before and incubated in LPDS media for 24 hours prior to incubation with the LDL-CTL-oleate ester for 2, 6, and 24 hours. Cells were washed with PBS and harvested by scraping in cold PBS. After centrifugation at 500g and 4° C for 3 min, the supernatant was removed and the sample was snap-frozen in liquid nitrogen. Frozen cell pellets were incubated for 5 minutes at 90°C with pre-heated 155 mM ammonium formate buffer and intermittent vortexing, then placed on ice. The samples were sonicated for 20 seconds at 40% power using a Branson tip sonicator. An aliquot of each cell lysate was used to determine the protein concentration using the BCA assay (ThermoFisher Scientific).

All samples, including aliquots of the LDL-CTL-oleate ester preparation, were subjected to MS^ALL^-based lipidomic analysis, as previously described [78–80]. In brief, the volume of each sample corresponding to 15 µg protein was mixed with 155 mM ammonium formate buffer up to 200 µL in total and spiked with 30 μL internal lipid standard mixture. From the LDL-CTL- oleate ester preparation, 1 and 10 nmol were used. Samples were subsequently extracted by adding 990 μL chloroform/methanol (2:1, v/v), vigorous shaking (1400 rpm, 90 min), followed by centrifugation (6000 g, 3 min) to promote phase separation. Lower organic phases were collected in new vials and vacuum evaporated, and stored at -20°C until analysis. Prior to lipidomic analysis, each lipid extract was dissolved in 100 μL chloroform/methanol (1:2, v/v), gently shaken (700 rpm, 3 min) and centrifuged (20,000 rpm, 3 min). Ten (10) μL lipid extracts were loaded in 96-well plates, mixed with 12.9 µL 13.3 mM ammonium formate in 2-propanol for analysis in positive ion mode.

Lipid extracts were analyzed by MS^ALL^ analysis in positive ion mode using an Orbitrap Fusion Tribrid (Thermo Fisher Scientific) equipped with a robotic nanoflow ion source, TriVersa NanoMate (Advion Biosciences). High-resolution survey Fourier transform MS (FTMS^1^) spectra were recorded across the range of m/z 280 to 2000 using a max injection time of 100 ms, automated gain control at 1e5, three microscans and a target resolution of 500,000. Consecutive FTMS^2^ spectra were acquired for all precursors in the range of m/z 398.3 to 1000.8 in steps of 1.0008 Da, recorded across a range starting from m/z 150 until the precursor m/z value + 10 Da, using max injection time of 100 ms, automated gain control at 5e4, a quadrupole ion isolation width of 1.0 Da, HCD fragmentation and a target resolution of 30,000. Targeted monitoring of ammoniated cholesterol (m/z 404.38869) and the internal standard cholesterol+2H7 (m/z 411.43265) was done by MSX analysis using max injection time of 600 ms, automated gain control at 5e4, five microscans, a target resolution of 120,000, HCD fragmentation at 8% and a quadrupole ion isolation width of 1.5 Da for each precursor [79, 80]. Two additional targeted MSX events were added, using the same parameters as above, to monitor ammoniated CTL (m/z 400.36) together with the cholesterol+2H7 internal standard, as well as ammoniated CTL 18:1 ester (m/z 664.60) together with the internal standard CE 16:1+2H7 (m/z 647.64), the latter pair at a collision energy of 12%.

Lipid molecules were identified using ALEX^123^ software, using an extended lipid fragmentation database covering the CTL 18:1 ester as well as its remodeled analogs with 14:0 to 22:6 fatty acyl chains, and quantified using a data processing pipeline in SAS 9.4 (SAS Institute) [78, 81–83]. [Note that although several endogenous CE species are isomeric with (re-esterified) CTL esters, the specific CTL fragment enables distinct quantification.]

