## Supplemental material for "Live-cell imaging and lipidomics of low density lipoprotein containing intrinsically fluorescent cholesteryl esters"

**Synthesis of cholestatrienol esters and ethers**

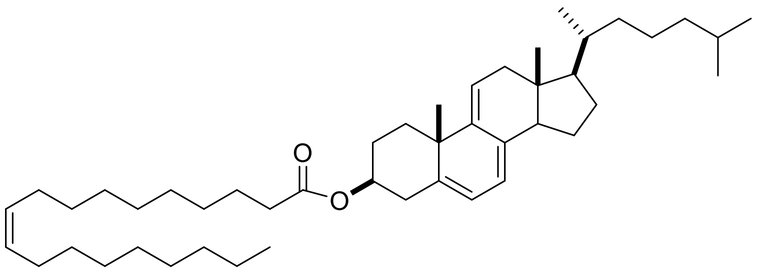

To a solution of cholestatrienol (100 mg, 0.262 mmol), oleic acid (89 mg, 0.314 mmol), and 4-dimethylaminopyridine (64 mg, 0.524 mmol) in methylene chloride (20 mL) was added *N*,*N'*-dicyclohexylcarbodiimide (65 mg, 0.314 mmol) in methylene chloride (2 mL) dropwise at 0 °C. The mixture was stirred at room temperature for 16 h. Aqueous $NH_4Cl$ was added and extracted with methylene chloride (100 mL x2). The solvent was removed, and the residue was purified by flash chromatography (silica gel, eluted with hexanes/EtOAc/$Et_3N$ 100/2/1, v/v/v) to give product (96 mg, 55%): $^1$H NMR (400 MHz, $CDCl_3$) δ 5.71-5.70 (m, 1H), 5.53-5.52 (m, 1H), 5.42-5.31 (m, 1H), 5.36-5.32 (m, 2H), 4.72-4.66 (m, 1H), 2.58-0.83 (m, 67H), 0.58 (s, 3H); $^{13}$C NMR (100 MHz, $CDCl_3$) δ 173.1, 144.0, 140.1, 135.6, 130.0, 129.7, 122.6, 119.0, 115.6, 73.7, 56.3, 50.9, 43.0, 42.2, 39.5, 39.3, 38.1, 37.5, 36.0, 35.9, 34.6, 31.9, 30.2, 29.8, 29.7, 29.5, 29.3 (3 x C), 29.2, 29.1 (2 x C), 28.5, 28.3, 28.0, 27.2, 27.1, 25.0, 23.9, 22.8, 22.7, 22.5, 18.4, 14.1, 11.4.

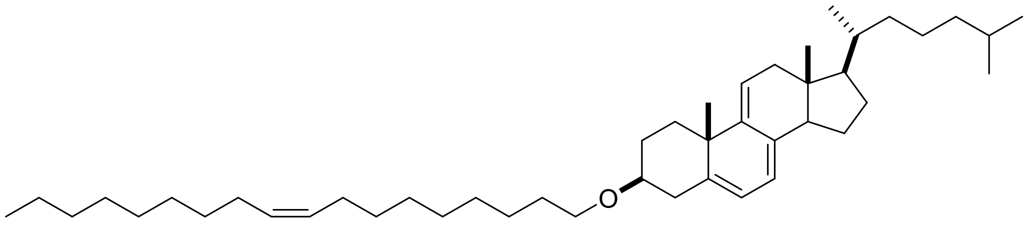

A mixture of 60% suspension of NaH in mineral oil(400 mg, 10 mmol), (Z)-1-iodooctadec-9-ene (907 mg, 2.4 mmol), and cholestatrienyl acetate (107 mg, 0.25 mmol) in THF (25 mL) was heated at reflux for 14 h. After cooling, water was slowly added and the mixture was extracted with EtOAc (100 mL x3). The solvent was removed under reduced pressure, and the residue was purified by flash chromatography (silica gel, eluted with hexanes/EtOAc/$Et_3N$ 100/1/1-100/2/1, v/v/v) to give product (152 mg, 96%): $^1$H NMR (400 MHz, $CDCl_3$) δ 5.69-5.68 (m, 1H), 5.52-5.51 (m, 1H), 5.42-

5.32 (m, 3H), 3.48-3.40 (m, 3H), 2.47-0.88 (m, 65H), 0.58 (s, 3H); $^{13}$C NMR (100 MHz, $CDCl_3$) δ 144.3, 141.8, 135.3, 129.9, 129.8, 122.3, 118.0, 115.6, 79.4, 71.0, 68.3, 56.4, 50.9, 43.0, 42.2, 39.7, 39.5, 38.5, 38.4, 36.0, 35.9, 31.9, 30.4, 30.2, 29.76, 29.74, 29.70, 29.6, 29.52, 29.49, 29.3, 29.24, 29.18, 29.1, 28.5, 28.0, 27.2, 26.2, 23.9, 22.8, 22.7, 22.5, 18.4, 14.1, 11.3.

**Kinetic modelling**

For analysis of the uptake and hydrolysis kinetics of LDL-associated CTL-ester, a sequential transport model was used. This model consists of three pools, the LDL containing CTL-ester in the medium (pool A), the LDL particles inside early and late endosomes after cellular uptake (pool B) and the liberated CTL upon hydrolysis of the LDL-derived CTL-ester in LE/LYSs (pool C; see inset in Fig. 3B, Fig. S5A and Fig. S6). The ordinary differential eqation (ODE) system for this model reads:

$$\frac{dA}{dt} = -k_1 \cdot A \qquad (1)$$

$$\frac{dB}{dt} = k_1 \cdot A - k_2 \cdot B \qquad (2)$$

$$\frac{dC}{dt} = k_2 \cdot B \qquad (3)$$

The solution of this ODE system can be found by several methods including spectral decomposition of the system matrix, Laplace transform and convolution method and reads for the initial condition $A(0) = A_0$, $B(0) = C(0\ ) = 0$:

$$A(t) = A_0 \cdot \exp(-k_1 \cdot t) \qquad (4)$$

$$B(t) = \frac{A_0 \cdot k_1}{k_2 - k_1} \cdot (\exp(-k_1 \cdot t) - \exp\,(-k_2 \cdot t)) \qquad (5)$$

$$C(t) = A_0 \cdot \left(\frac{k_1}{k_2 - k_1} \cdot \exp(-k_2 \cdot t) - \frac{k_2}{k_2 - k_1} \cdot \exp(-k_1 \cdot t) + 1\right) \qquad (6)$$

The solution for $C(t)$ of Eq. 6 was fit to the experimental data.

For analysis of cellular uptake of LDL-associated CEs and TAG species, a 2-pool model was used, which is identical to the 3-pool model without pool C. Its ODE system is equal to that of Eqs. 1 and 2 for $k_1 = k > 0$ and $k_2 = 0$. Its solution for pool A is the same as Eq. 4, while that for pool B reads:

$$B(t) = A_0 \cdot (1 - \exp(-k \cdot t)) \qquad (7)$$

Eq. 7 was used in the hierarchical Bayesian model as well as in analysis of CTL reesterification.

**Single particle tracking and quantification of interorganelle distances**

Upon thresholding the time-lapse images or 3D image stacks of the ER markers, the resulting binary image sequence was skeletonized, and an Euclidian distance transform was calculated in physical dimensions (µm). For the 3D data, the 3D Euclidian distance transform implemented in ImageJ was used. For 3D data, the endo-lysosomes were segmented using the Tango plugin to ImageJ, as described [1, 2]. For time-lapse data, positions of endo-lysosomes containing Alexa488-LDL were determined by single particle tracking, as described [2]. An anomalous diffusion model was fit to the calculated mean square displacement of individual trajectories [2].

**Table S1. Phosphocholine and -ethanolamine glycerol ester composition of reconstituted LDL.** Relative abundance of phosphatidylethanolamine (PE) and phosphatidylcholine (PC).

| PE species | Concentration (mol% of all lipids) | PC species | Concentration (mol% of all lipids) |
|---|---|---|---|
| PE 32:1 | 0.0096 | PC 30:1 | 0.0164 |
| PE 34:1 | 0.0014 | PC 30:0 | 0.0015 |
| PE 36:1 | 0.0084 | PC 32:2 | 0.0072 |
| PE 36:2 | 0.0174 | PC 32:1 | 0.0113 |
| PE 36:4 | 0.0006 | PC 32:0 | 0.0168 |
| PE 36:5 | 0.0006 | PC 34:4 | 0.0001 |
| PE 38:1 | 0.0043 | PC 34:3 | 0.0092 |
| PE 38:2 | 0.0193 | PC 34:2 | 0.6686 |
| PE 38:4 | 0.0013 | PC 34:1 | 0.2758 |
| PE 38:5 | 0.0003 | PC 34:0 | 0.0054 |
| PE 40:3 | 0.0025 | PC 36:5 | 0.0131 |
| PE 40:4 | 0.0016 | PC 36:4 | 0.2405 |
| PE40:6 | 0.0004 | PC 36:3 | 0.1711 |
| PE42:6 | 0.0008 | PC 36:2 | 0.3615 |
| | | PC 36:1 | 0.0409 |
| | | PC 36:0 | 0.0002 |
| | | PC 38:7 | 0.0022 |
| | | PC 38:6 | 0.0517 |
| | | PC 38:5 | 0.0484 |
| | | PC 38:4 | 0.1308 |
| | | PC 38:3 | 0.0460 |
| | | PC 38:2 | 0.0026 |
| | | PC 40:8 | 0.0006 |
| | | PC 40:7 | 0.0023 |
| | | PC 40:6 | 0.0195 |
| | | PC 40:5 | 0.0085 |
| | | PC 40:4 | 0.0024 |
| | | PC 44:9 | 0.0002 |
| | | PC 44:4 | 0.0002 |

**Table S2. Sphingolipid composition of reconstituted LDL.** Relative abundance of ceramide and sphingomyelin species were determined by Lipid-MS after lipid extraction from LDL particles reconstituted with CTL-oleate ester.

| Ceramide species | Concentration (mol% of all lipids ) | Sphingomyelin species | Concentration (mol% of all lipid ) |
|---|---|---|---|
| Cer 32:2,2 | 0.0011 | SM 32:1,2 | 0.0190 |
| Cer 32:2,3 | 0.0011 | SM 34:2,2 | 0.0316 |
| Cer 34:0,3 | 0.0029 | SM 34:1,2 | 0.2885 |
| Cer 36:0,2 | 0.0005 | SM 34:0,2 | 0.0103 |
| Cer 36:2,3 | 0.0019 | SM 36:2,2 | 0.0201 |
| Cer 36:0,3 | 0.0053 | SM 36:1,2 | 0.0475 |
| Cer 38:1,3 | 0.0016 | SM 36:0,2 | 0.0016 |
| Cer38:0,3 | 0.0000 | SM 38:2,2 | 0.0126 |
| Cer 40:0,2 | 0.0010 | SM 38:1,2 | 0.0414 |
| Cer 42:2,2 | 0.0025 | SM 38:2,3 | 0.0001 |
| Cer 42:1,2 | 0.0219 | SM 40:2,2 | 0.0509 |
| Cer 42:0,2 | 0.0017 | SM 40:1,2 | 0.0774 |
| Cer 42:2,3 | 0.0007 | SM 40:2,3 | 0.0002 |
| Cer 46:0,3 | 0.0021 | SM 42:2,2 | 0.1268 |
| | | SM 42:1,2 | 0.0448 |
| | | SM 42:2,3 | 0.0002 |
| | | SM 42:1,3 | 0.0002 |
| | | SM 44:2,2 | 0.0003 |

**Table S3. Kinetic parameters of CTL-ester transport and hydrolysis.** Kinetic parameters are shown for the sequential model as determined by non-linear regression to the data of the indicated experiment. Results of the Bayesian regression are shown for comparison. The rate constant and half-time for the reesterification is also given.

| Experiment | Start amount in pool $A_0$ | Rate constant for first step, $k_1$ | Rate constant for second step, $k_2$ | Rate constant for re-esterification | Technique /Figure |
|---|---|---|---|---|---|
| CTL-ester uptake & transport to LE/LYSs | 96.49 (a.u.) | 0.0096 min$^{-1}$ ($t_{1/2}$ = 72.2 min) | 0.0096 min$^{-1}$ ($t_{1/2}$ = 72.2 min) | -- | Imaging/ Figure 3B (orange) |
| CTL-ester uptake & transport to LE/LYS (Bayesian fit)s | 96.602 ± 0.913 | 0.0096 ± 0.002 min$^{-1}$ ($t_{1/2}$ = 72.2 min) | 0.0096 ± 0.003 min$^{-1}$ ($t_{1/2}$ = 72.2 min) | -- | Imaging/ Figure S5 |
| Hydrolysis of CTL-oleate ester (all free) | 0.354 pmol/µg protein | 0.0054 min$^{-1}$ ($t_{1/2}$ = 129.2 min) | 0.0054 min$^{-1}$ ($t_{1/2}$ = 129.2 min) | -- | Lipid-MS/ Figure 4A (orange) |
| Hydrolysis of CTL-oleate ester ($k_1$ = *fixed*) | 0.355 pmol/µg protein | *0.0096 min$^{-1}$ ($t_{1/2}$ = 72.2 min)* | 0.0035 min$^{-1}$ ($t_{1/2}$ = 198.0 min) | -- | Lipid-MS/ Figure 4A (green) |

| | | | | | |
|---|---|---|---|---|---|
| Formation of CTL-ester (18:0) | -- | -- | -- | $k$ = 0.0045 $min^{-1}$ ($t_{1/2}$ = 152.7 min) | |
| Hydrolysis of CTL-oleate ester (Bayesian fit) | 0.333 ± 0.044 pmol/µg protein | 0.0096 ± 0.0015 $min^{-1}$ ($t_{1/2}$ = 72.2 min) | 0.0070 ± 0.0019 $min^{-1}$ ($t_{1/2}$ = 99.02 min) | -- | Lipid-MS/ Figure S5 |

**Table S4. Kinetic parameters of cellular uptake of LDL-derived CEs.** Kinetic parameters are shown for the mono-exponential model as implemented in a hierarchic Bayesian framework applied to the data of the indicated experiment. Amplitudes were determined separately, while the rate constant was shared for each data set. Values are given as mean ± standard deviation of the posterior distribution. See Figure S8 for time courses and Monte Carlo simulations.

| **CE species** | **Start amount in pool $A_0$** | **Rate constant for cellular uptake, *k*** | **Half-time for cellular uptake, $t_{1/2}$** |
|---|---|---|---|
| 16:0 | 1.273 ± 0.038 pmol/µg protein | 0.00358 ± 0.00003 $min^{-1}$ | 193.4 min |
| 16:1 | 0.439 ± 0.037 pmol/µg protein | | |
| 18:1 | 10.191 ± 0.043 pmol/µg protein | | |
| 18:2 | 10.207 ± 0.042 pmol/µg protein | | |

**Table S5 Kinetic parameters of cellular uptake of LDL-derived TAGs.** Kinetic parameters are shown for the mono-exponential model as implemented in a hierarchic Bayesian framework applied to the data of the indicated experiment. Amplitudes were determined separately, while the rate constant was shared for each data set. Values are given as mean ± standard deviation of the posterior distribution. See Figure S9 for species abundance and time courses.

| **TAG species** | **Start amount in pool $A_0$** | **Rate constant for cellular uptake, *k*** | **Half-time for cellular uptake, $t_{1/2}$** |
|---|---|---|---|
| TAG50:1 | 0.605 ± 0.044 pmol/µg protein | 0.00168 ± 0.00008 $min^{-1}$ | 411.8 min |
| TAG50:2 | 0.509 ± 0.044 pmol/µg protein | | |
| TAG50:3 | 0.214 ± 0.044 pmol/µg protein | | |
| TAG52:2 | 1.404 ± 0.047 pmol/µg protein | | |
| TAG52:3 | 1.516 ± 0.048 pmol/µg protein | | |
| TAG52:4 | 0.994 ± 0.046 pmol/µg protein | | |

| | |
|---|---|
| TAG54:3 | 0.913 ± 0.046 pmol/µg protein |
| TAG54:4 | 1.978 ± 0.054 pmol/µg protein |

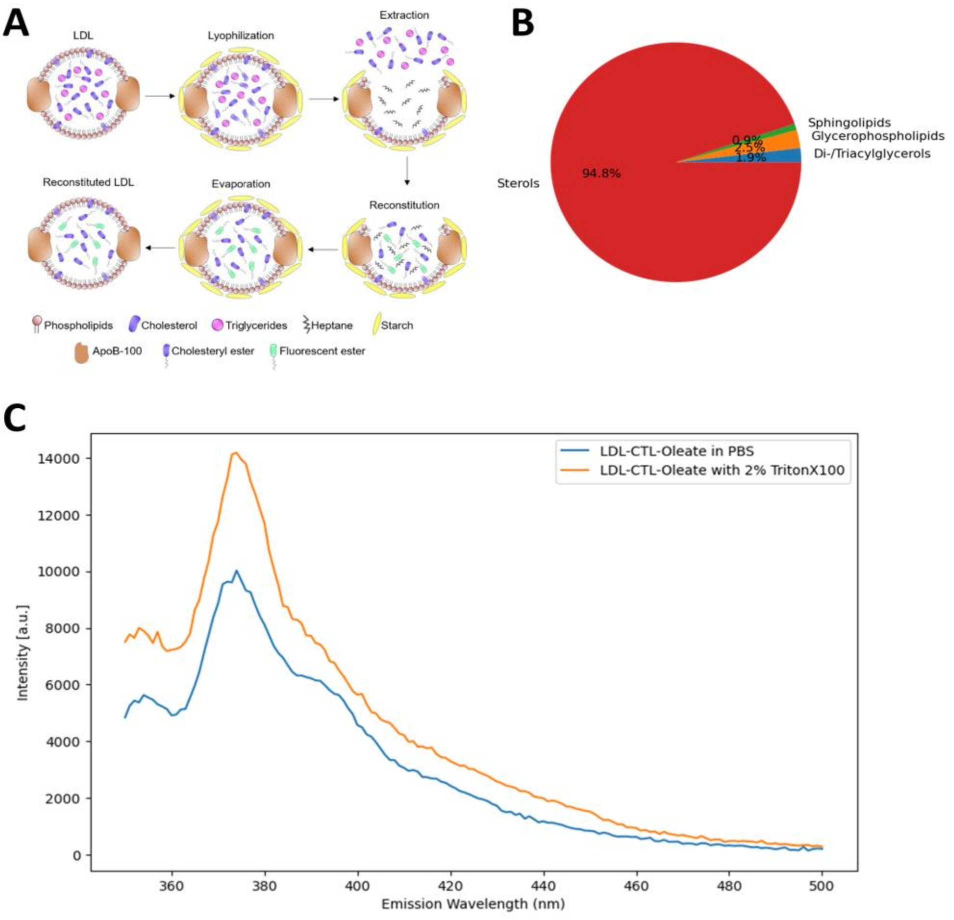

**Figure S1. Characterization of reconstituted LDL particles.** A, scheme of the reconstitution procedure, B, lipid compsotion of the LDL particles containing fluorescent CTL-oleate esters, C, emission spectra of the particles in the absence (blue line) or presence of the detergent Triton X-100 (orange line). Excitation was set to 328 nm, and 10 µl of LDL-CTL-oleate in 500 µl PBS were recorded. See main text for further details.

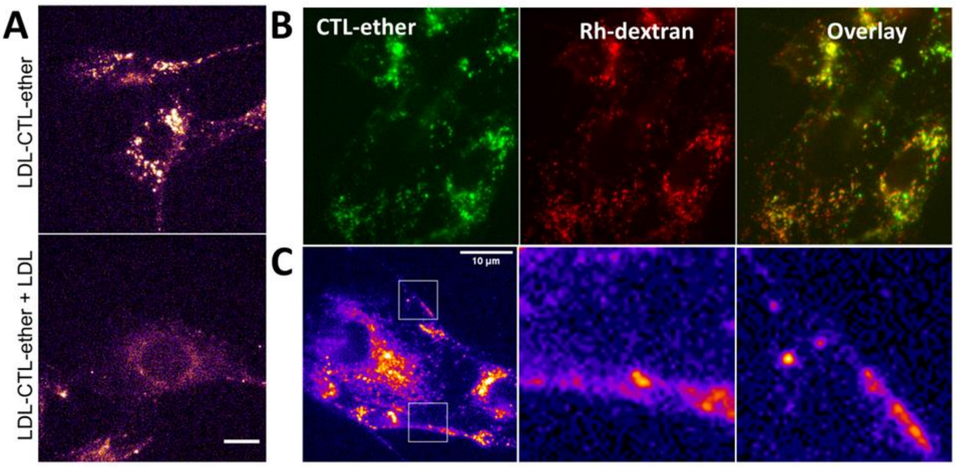

**Figure S2. Live-cell imaging and Lipid-MS of LDL containing CTL-oleate ether.** A, fibroblasts were incubated in LPDS medium overnight before adding 0.1 mg/ml LDL with CTL-oleate ether for 24h (A, upper panel). In a separate dish, excess unlabeled LDL (4 mg/ml) was added during the incubation with reconstituted LDL particles (lower panel in A). B, to stain LE/LYSs 0.5 mg/ml Rh-dextran was added together with labeled LDL particles, showing that the CTL-ether accumulates in endo-lysosomes. C, cells were incubated as in A (upper panel), and zoomed boxes illustrate the patchy signal in the PM stemming from labeled LDL particles bound to their receptor on the cell surface.

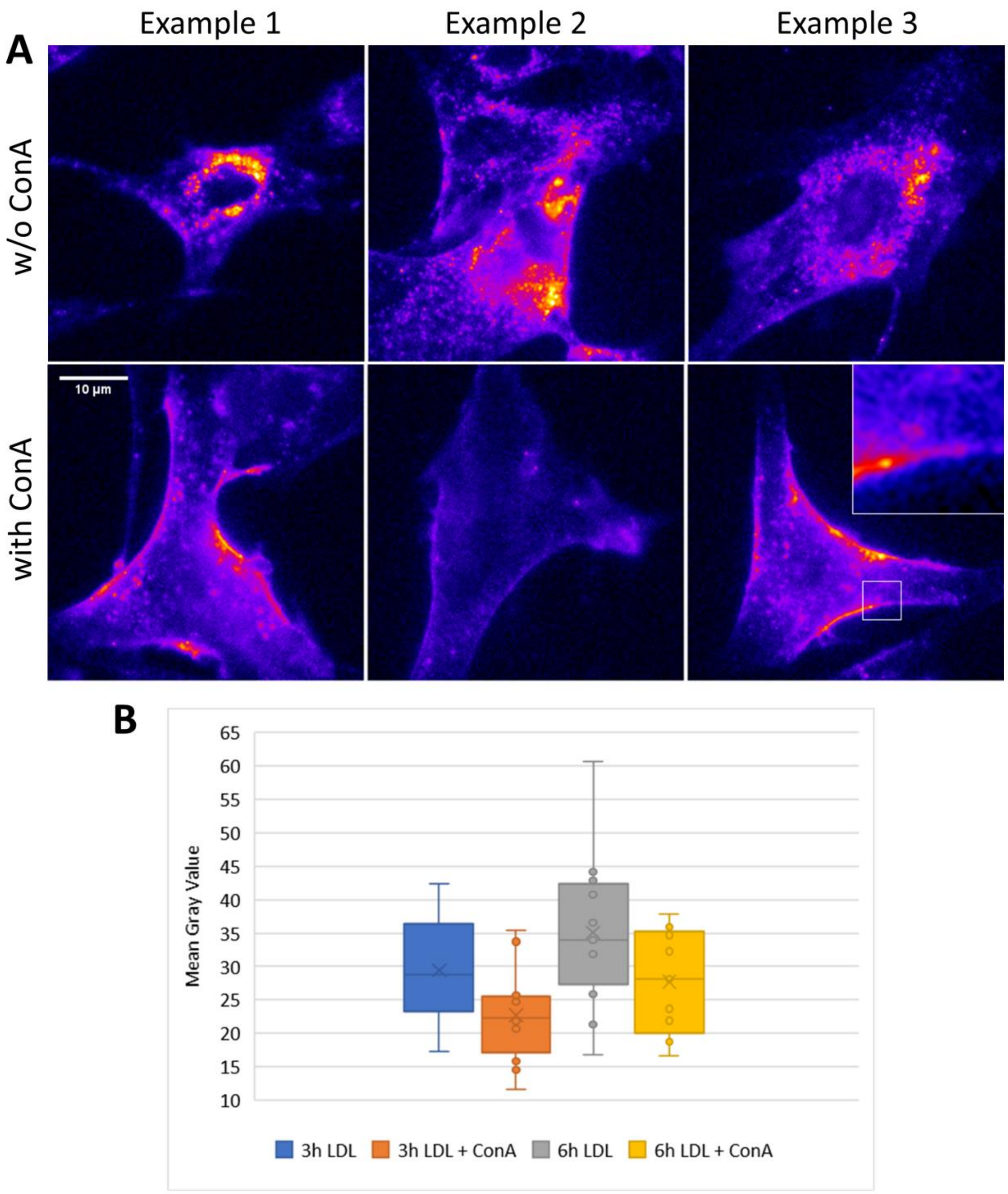

**Figure S3. Concanamycin A blocks uptake of reconstituted LDL particles.** Human fibroblasts were incubated overnight in LPDS medium with or without 250 nM ConA. On the day of microscopy, the cells were loaded with LDL-CTL-ester for 6 h. A, three image examples, which are equally scaled, are shown with a FIRE LUT with high intensities yellow and low intensities in blue. B, quantification of cell-associated CTL(ester) fluorescence expressed as mean +/- S.D. of 15 cells from one experiment.

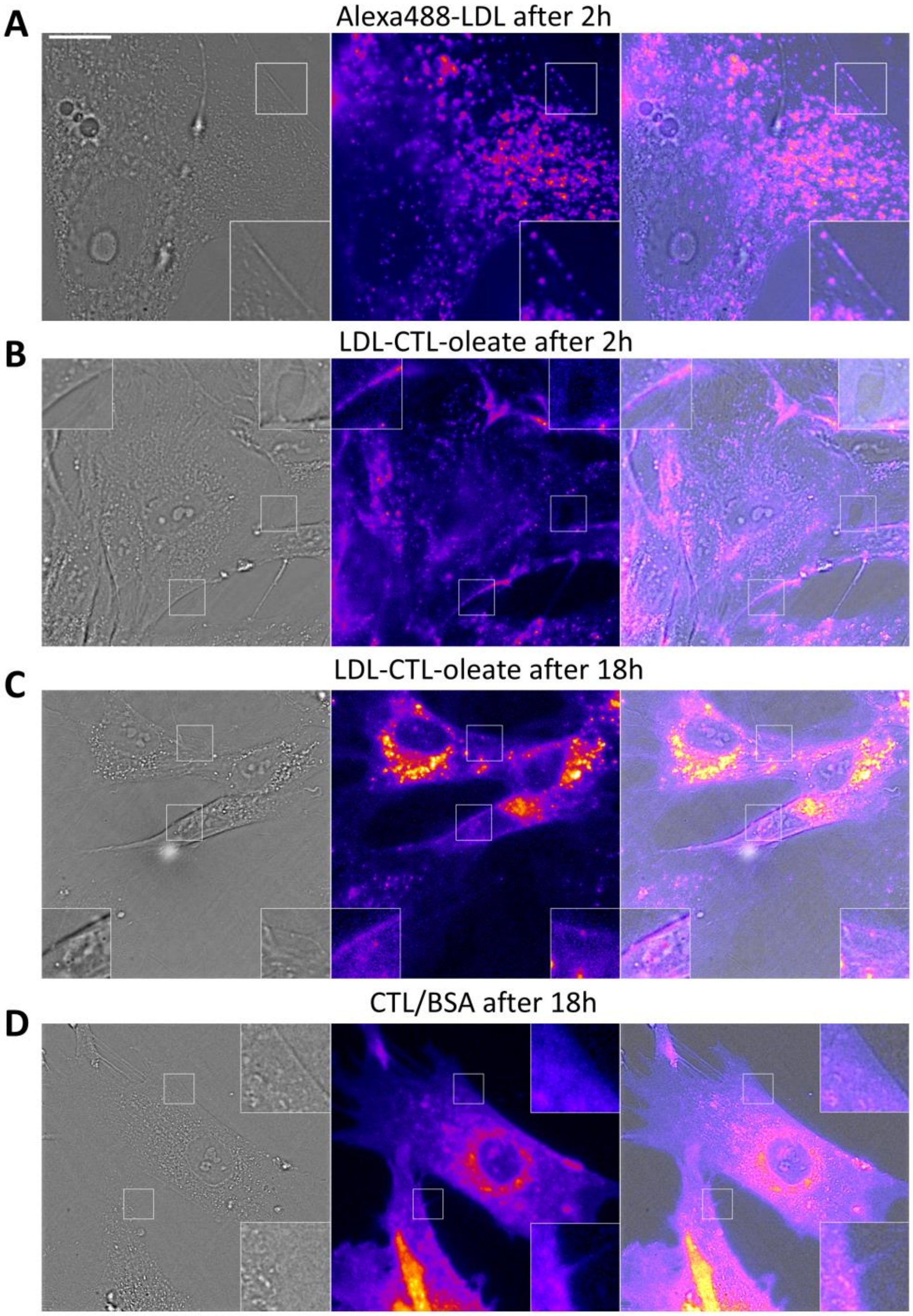

**Figure S4. Comparison of plasma membrane fluorescence of LDL particles.** A-C, human fibroblasts were incubated in LPDS medium overnightA, The next day, they were labeled with 0.1 mg/ml Alexa488-LDL (A) or LDL-CTL-oleate for either 2h (B) or 24h (C). D, for comparison cells were labeled in an independent experiment with CTL loaded onto BSA ovrnight, which results in an intense staining of the PM (see inset). Insets in A-C show the fluorescence of Alexa488 (A) and CTL(ester) (B and C) or CTL (D) in the PM. Bar, 10 µm.

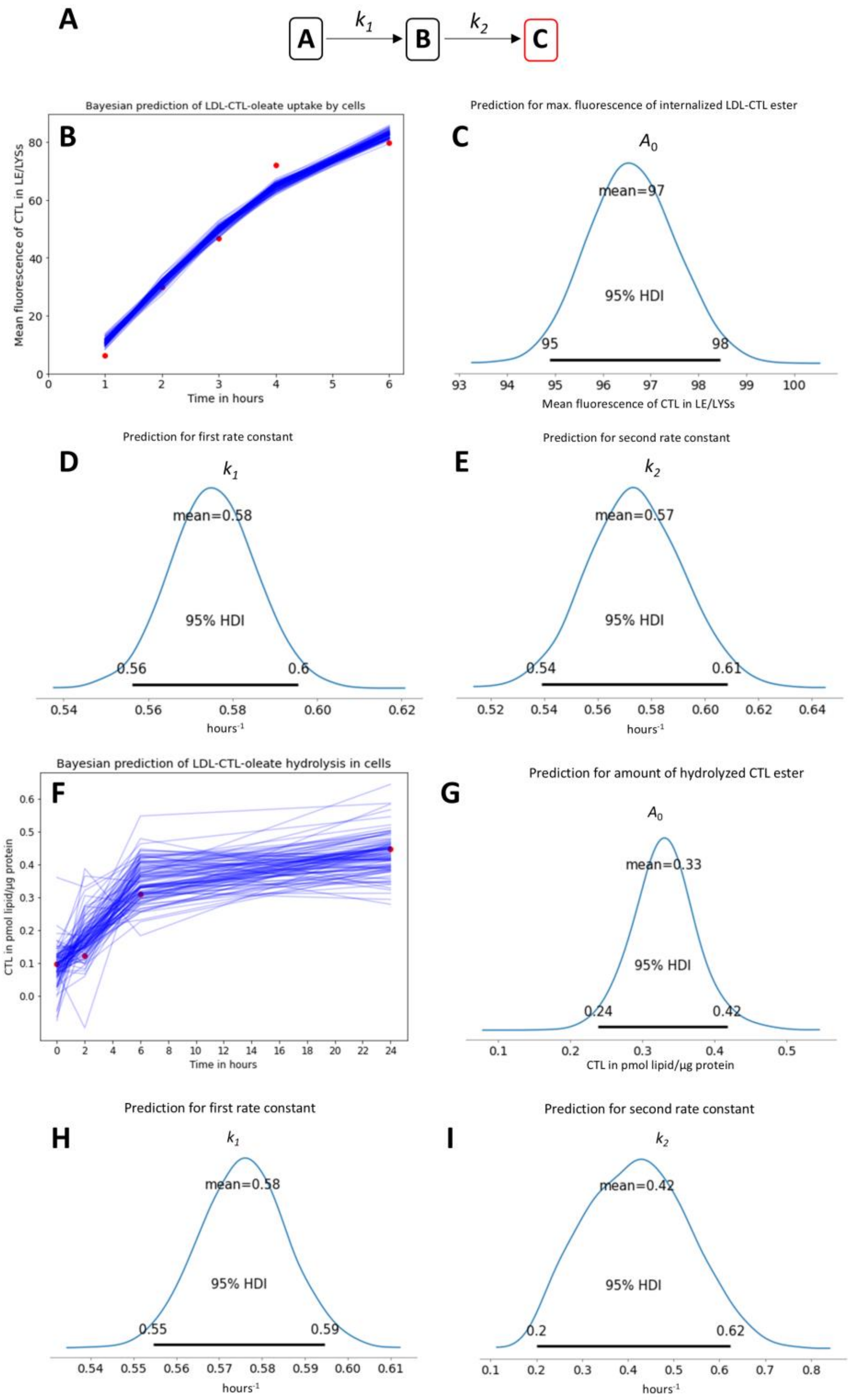

**Figure S5. Bayesian analysis of transport and metabolism of LDL-derived CTL-esters.** The sequential model shown in A was fit to the kinetics of fluorescence increase in LE/LYSs (B-E) and separately to the hydrolysis kinetics of CTL-ester, determined by Lipid-MS (F-I); compare with Figure 4A and main text. B and F show the mean of the data (red dots) overlayed with 100 simulated time courses sampled from the model posterior (blue lines). From the latter distribution the inferred values for the amplitudes (C and G) and rate constants $k_1$ (D and H) and $k_2$ (E and I) are shown with the 95% highest density interval (HDI) indicated. The posterior for each parameter was determined by Monte Carlo sampling in PyMC3 including the mean value in hours$^{-1}$ from four separate runs with 2000 steps each.

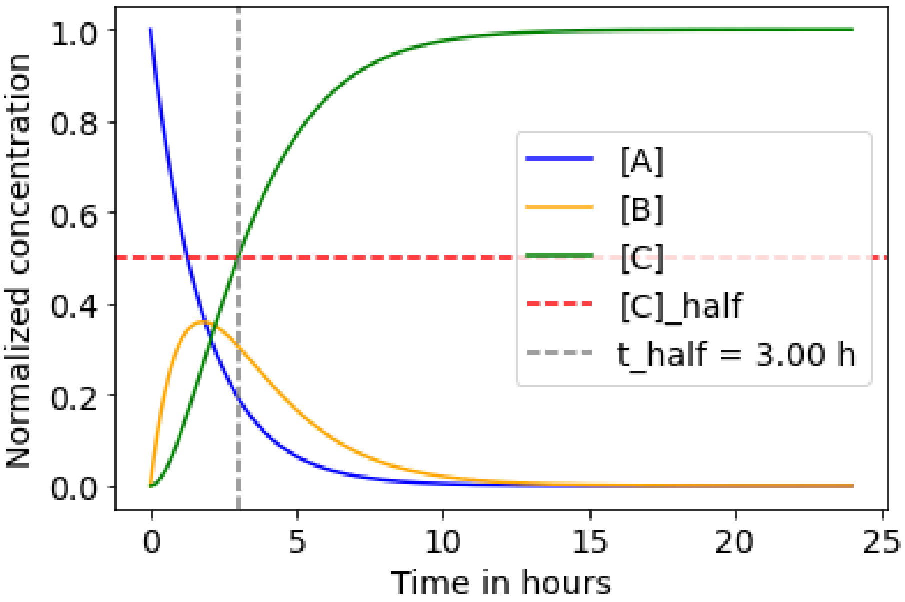

**Figure S6. Simulation of sequential kinetic model.** All three compartments were simulated with the rate constants $k_1$ and $k_2$ shared between the two data sets of Figure 3B and Figure 4A for inference with the hierarchical Bayesian model. The initial amount of LDL in the medium (poolA in blue) is set to 1, while the other two pools were initially zero The time course for LDL in early and late endosomes is shown in yellow (pool B) and that for LDL hydrolysis in endo-lysosomes in green (pool C). The half-time of the entire process was calculated as half-maximal value of poolC (red dahsed line) with the perpendicular grey dashed line refering to the corresponding half-time (t_half) of 3 hours. See main text for further details.

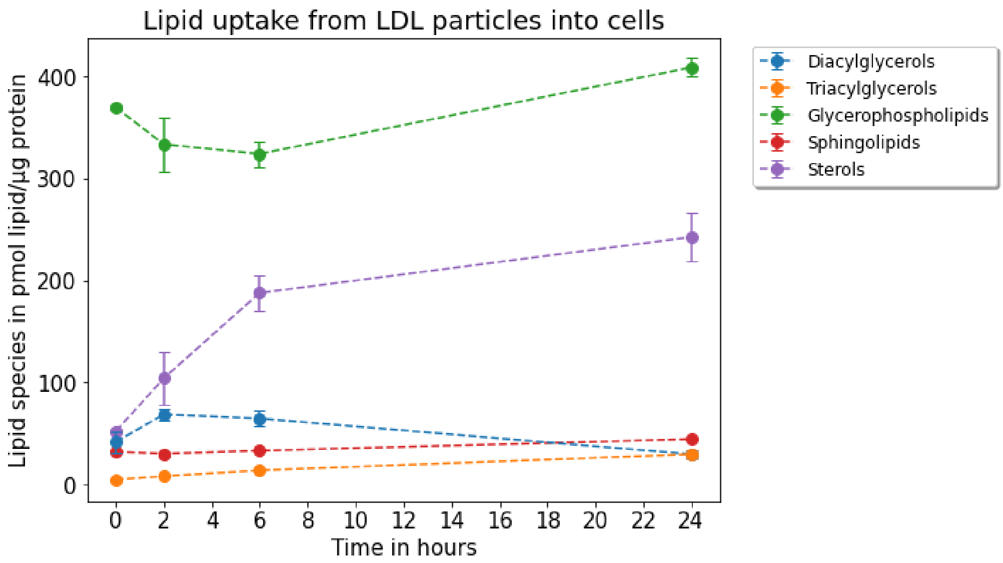

**Figure S7. Lipid-MS of cell-associated lipids classes.** Fibroblasts were incubated in LPDS medium containing 0.1 mg/mL LDL with CTL-oleate ester for the indicated times. Plots show the quantification of cell-associated sterols (purple), sphingolipids (red), diacylglycerols (blue), triacyclglycerols (orange), and glycerophospholipids (green). Data is shown as mean +/- SEM of three technical replicates.

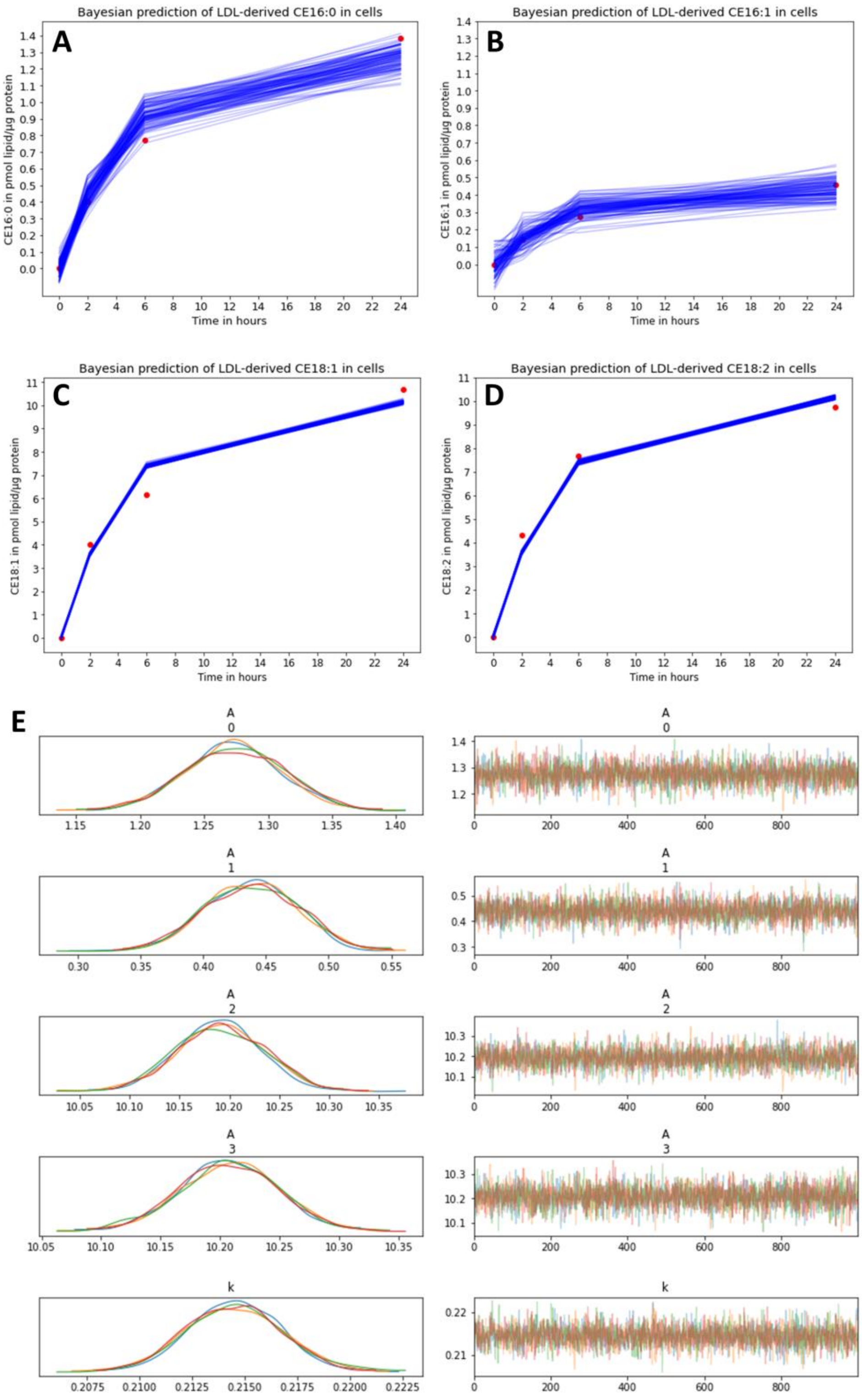

**Figure S8. Bayesian modeling of LDL-associated CEs.** CEs found to be specifically enriched in LDL were quantified over time (A-D, mean values shown as red dots) and analyzed using the hierarchical mono-exponential Bayesian model with simulations sampled using the posterior parameter estimates shown as blue lines in A-D. The analysis included CE16:0 (A), CE16:1 (B), CE18:1 (C) and CE18:2 (D). E shows the result of a Monte Carlo simulation of the model with the posteriors for each parameter shown in the left panels and the simulated traces in the right panels. Infered parameters were the four amplitudes (named A0 to A3 in E) and the rate constant k. The colors correspond to four independent runs of the model with 1000 steps each. All four estimates coincide well, showing the reproducibility of the analysis.

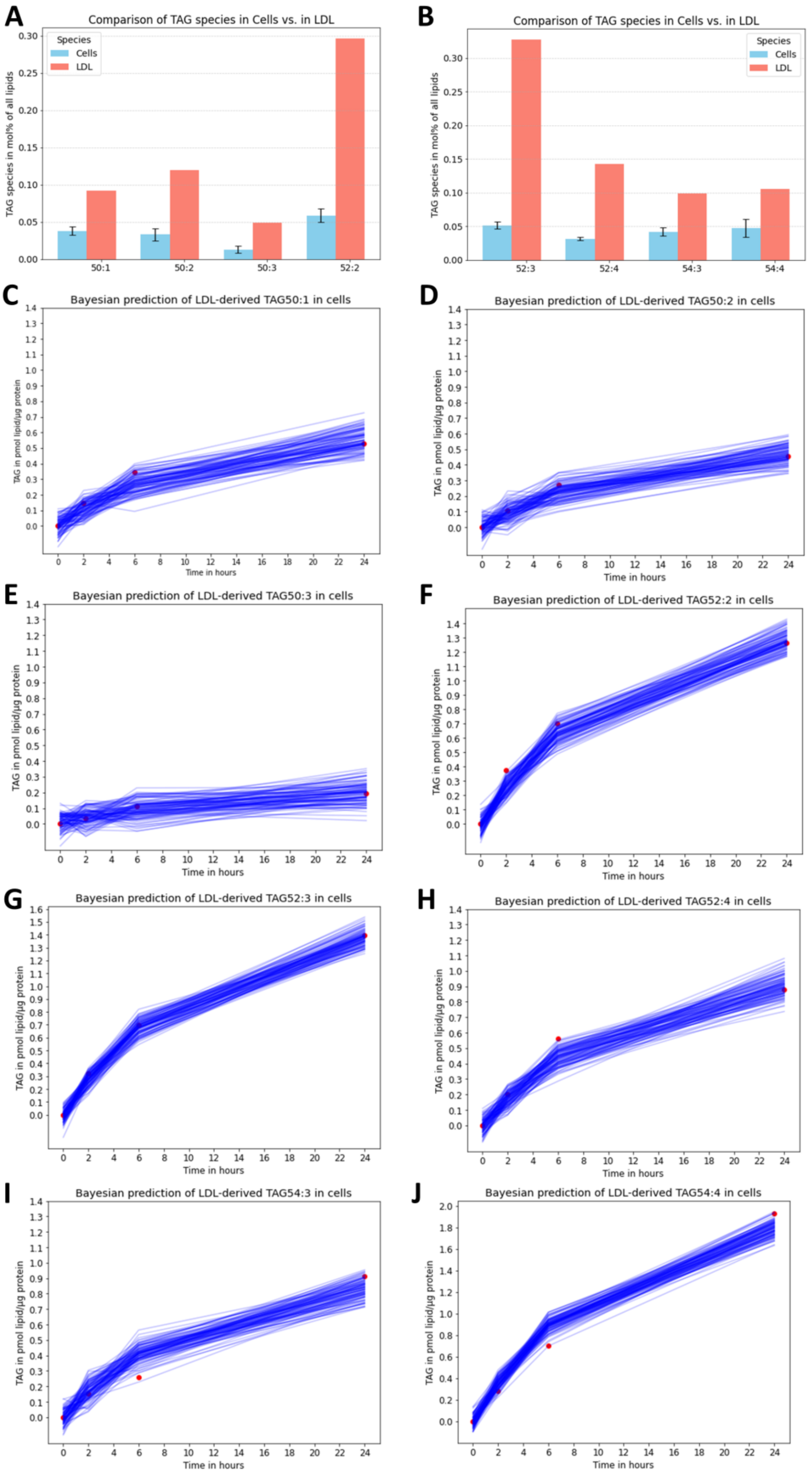

**Figure S9. Bayesian modeling of LDL-associated TAGs.** Selected TAGs were quantified in non-treated cells (A, B, blue bars) and in the reconstituted LDL particles (A, B, red bars). Their abundance in cells incubated with LDL was quantified over time (C-J, mean values shown as red dots) and analyzed using the hierarchical mono-exponential Bayesian model with simulations sampled using the posterior parameter estimates shown as blue lines in C-J. The analysis included TAG50:1 (C), TAG50:2 (D),. TAG50:3 (E), TAG52:2 (F), TAG52:3 (G), TAG52:4 (H), TAG54:3 (I), TAG54:4 (J). The simulated time courses were generated by Monte Carlo sampling from the posterior parameter distribution. Mean values of inferred parameters are shown in Tab. S3.

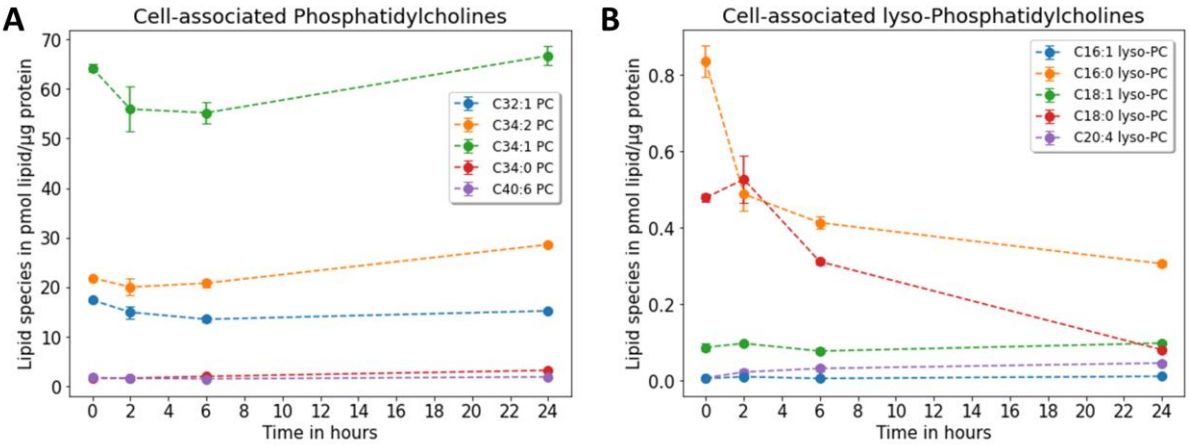

**Figure S10. Lipid-MS of cell-associated phosphatidylcholines in response to LDL loading.** Human fibroblasts were incubated in LPDS medium overnight, followed by incubation with 0.1 mg/ml LDL containing CTL-oleate-ester for the indicated times. Cells were washed, harvested and lipids extracted for analysis by Lipid-MS as described in Materials and methods. Time courses for cell-associated PC species are shown in pmol/µg protein for diacyl-PC esters (A) and monoacyl-PC esters (B). Data is shown as mean +/- SEM of three technical replicates.

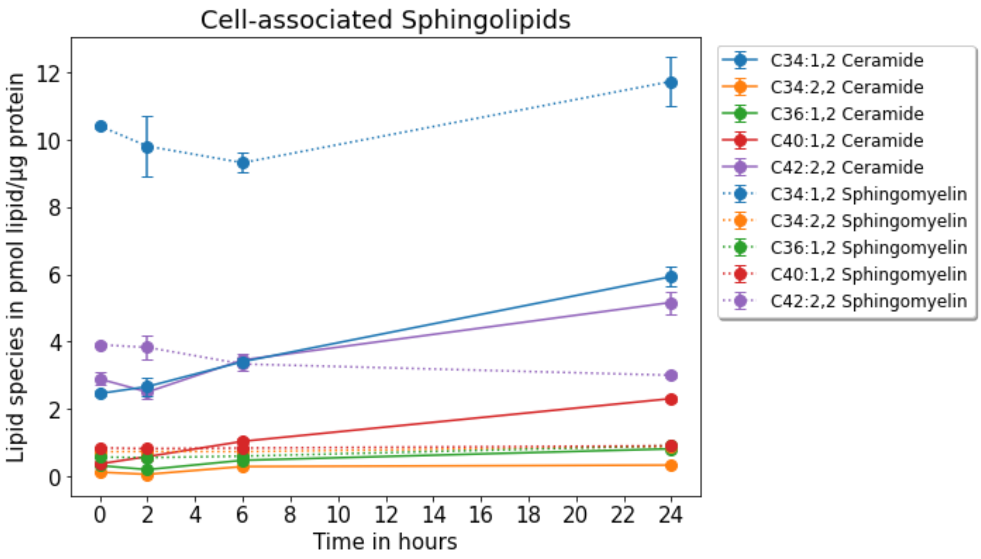

**Figure S11. Lipid-MS of cell-associated sphingolipids in response to LDL loading.** Human fibroblasts were treated as described in legend to Figure S10, above. Cells were washed, harvested and lipids extracted for analysis by Lipid-MS as described in Materials and methods. Time courses for selected cell-associated sphingolipids in pmol/µg protein is shown as straight lines for ceramides and dashed lines for SM species, respectively. Data is shown as mean +/- SEM of three technical replicates.

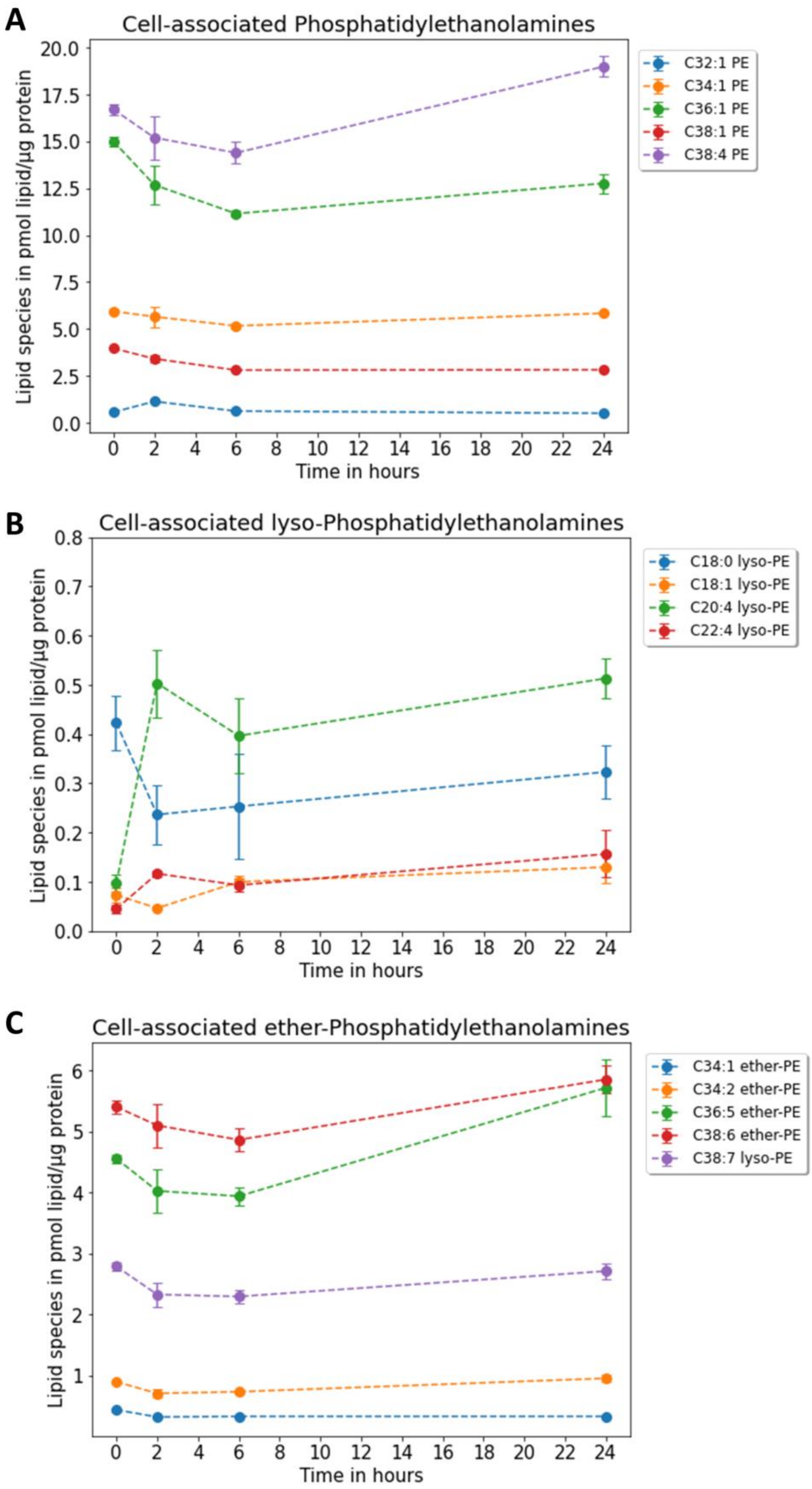

**Figure S12. Lipid-MS of cell-associated phosphatidyl-ethanolamines in response to LDL loading.** Human fibroblasts were incubated in LPDS medium overnight, followed by incubation with 0.1 mg/ml LDL containing CTL-oleate-ester for the indicated times. Cells were washed, harvested and lipids extracted for analysis by Lipid-MS as described in Materials and methods. Time courses for cell-associated PE species are shown in pmol/µg protein for diacyl-PE esters (A), monoacyl-PE esters (B) and diacyl-PE ethers (C). Data is shown as mean +/- SEM of three technical replicates.

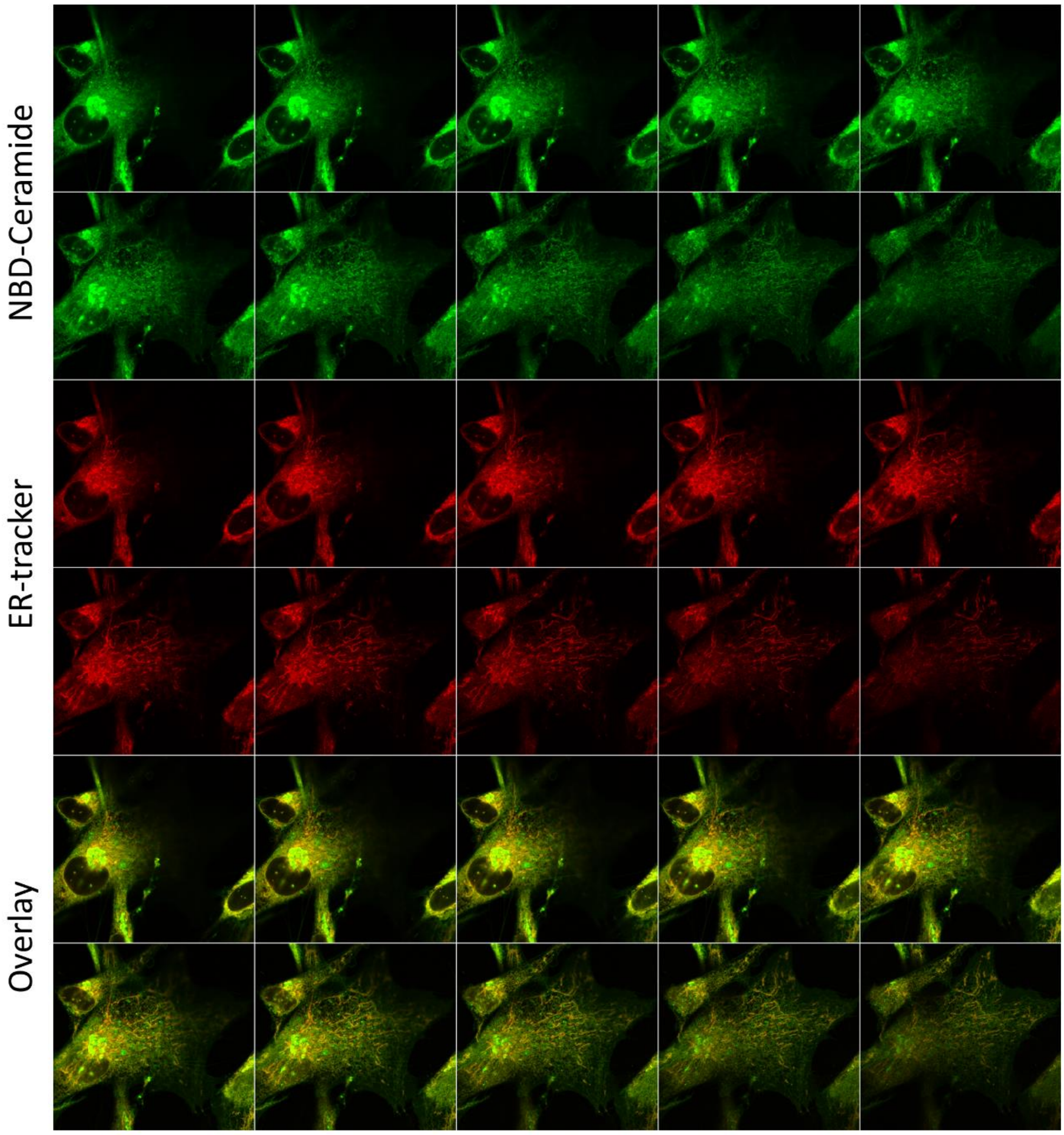

**Figure S13. Co-localization of NBD-Ceramide with ER Tracker.** Human fibroblasts were co-stained with 5 µM of NBD-Ceramide (upper panel) and ER-tracker Red (middle panel) and imaged on a confocal microscope. The lower panel is the color overlay, and successive images of a 3D stack 250 nm apart through the cell are shown.

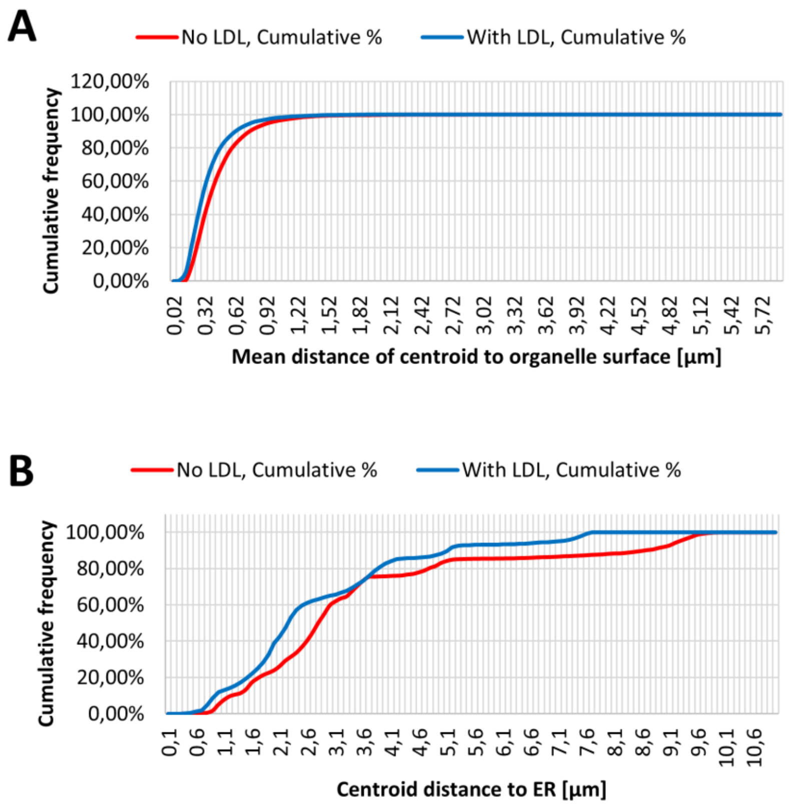

**Figure S14. Cumulative frequency plot of size of LE/LYSs and their distance to the ER.** Human fibroblasts were co-stained with MagicRed and NBD-Ceramide, as described in legend to Figure 6. 3D image stacks were acquired at a spinning disk confocal microscope, deconvolved and further analyzed to determine the distance between MagicRed stained LE/LYSs and NBD-Ceramide-labeled ER tubules (see Materials and methods). Cumulative frequency of endo-lysosome size (A) and distance to the ER (B) for cells incubated with LDL (blue) or without LDL (red) is shown.

The mean distance between the particle centroid and the organelle surface is a measure of endo-lysosome-size, while the shortest distance between endo-lysosome centroid and next ER tubule quantifies ER-LE/LYSs distance.

[1] J. Ollion, J. Cochennec, F. Loll, C. Escude, T. Boudier, TANGO: a generic tool for high-throughput 3D image analysis for studying nuclear organization, Bioinformatics, 29 (2013) 1840-1841.
[2] A.D. Juhl, C.W. Heegaard, S. Werner, G. Schneider, K. Krishnan, D.F. Covey, D. Wüstner, Quantitative imaging of membrane contact sites for sterol transfer between endo-lysosomes and mitochondria in living cells, Sci Rep, 11 (2021) 8927.
